# Molecular insights into substrate recognition and discrimination by the N-terminal domain of Lon AAA+ protease

**DOI:** 10.1101/2020.12.17.423354

**Authors:** Shiou-Ru Tzeng, Yin-Chu Tseng, Chien-Chu Lin, Chia-Ying Hsu, Shing-Jong Huang, Yi-Ting Kuo, Chung-I Chang

## Abstract

The Lon AAA+ proteases (LonA) is a ubiquitous ATP-dependent proteolytic machine, which selectively degrades damaged proteins or native proteins carrying exposed motifs (degrons). Here we characterize the structural basis for substrate recognition and discrimination by the N-terminal domain (NTD) of LonA. The results reveal that the six NTDs are attached to the hexameric LonA chamber by flexible linkers such that the formers tumble independently of the latter. Further spectral analyses show that the NTD selectively interacts with unfolded proteins, protein aggregates, and degron-tagged proteins by two hydrophobic patches of its N-lobe, but not intrinsically disordered substrate, α-casein. Moreover, the NTD selectively binds to protein substrates when they are thermally induced to adopt unfolded conformations. Collectively, our findings demonstrate that NTDs enable LonA to perform protein quality control to selectively degrade proteins in damaged states and suggest that substrate discrimination and selective degradation by LonA are mediated by multiple NTD interactions.

**Impact Statement:** The N-terminal domains enable Lon protease to discriminate and capture selected protein species for degradation by exposed hydrophobic patches and flexible linkages to the hexameric core complex.

## Introduction

The Lon AAA+ protease (LonA), previously known as the protease La, is an ATP-dependent protease distributed in prokaryote and eukaryote^1^. It forms a homo-hexamer to execute its biological function^2,3^. LonA belongs to the AAA+ (ATPases associated with various cellular activities) superfamily and contains an N-terminal domain (NTD), a middle ATPase domain with conserved Walker motifs for ATP hydrolysis, and a C-terminal protease domain (CTD) with a serine-lysine catalytic dyad in the active site^4,5^. LonA is responsible for degrading damaged or unfolded proteins, as well as native proteins bearing specific recognition elements, known as degradation tag or degrons^6,7^. How can LonA discriminate its substrate from other non-substrate proteins in a cell? Substrate recognition of LonA is thought to be mediated by its NTD^8–14^. The substrates are unfolded and translocated in an ATP-dependent process into a secluded chamber formed by the ATPase domains and the protease domains. Finally, the unfolded substrates inside the chamber are degraded into small peptide fragments^15,16^. The ATP-dependent translocation process of substrates into the hexameric chamber, formed by the AAA+ and protease domains, has been well characterized^17,18^. Structures of different N-terminal fragments of LonA from three species have been reported^19–22^, including *Escherichia coli* (EcLon), *Mycobacterium avium complex* (MacLon) and *Bacillus subtilis* (BsLon). However, no structural study has been conducted to analyze substrate interaction by the NTD either in isolation or in the context of full-length LonA.

Here we address the question of substrate recognition and discrimination by the NTD domains of LonA using nuclear magnetic resonance (NMR) spectroscopy. To understand how LonA selectively recognizes protein substrates being in damaged states, we have used a thermal stable LonA isolated from *Meiothermus taiwanensis* (termed MtaLonA henceforth), which allows temperature cycling experiments to be applied to change the populations of a protein substrate in folded and unfolded states. In this work, five various protein substrates have been employed for NMR experiments, which include: (i) Ig2 (domains 5 and 6 of the gelation factor from *Dictyostelium discoideum^23^*, hereafter abbreviated as Ig2) with an immunoglobulin (Ig) fold forming a dimer that is natively folded up to 40 °C; (ii) α-casein, which is an intrinsically disordered protein with no tertiary structure; (iii) native lysozyme with four disulfide bridges and reduced lysozyme forming loose and flexible aggregates; (iv) Ig2D5 (domain 5 of the gelation factor from *Dictyostelium discoideum*), which is used for characterization of substrate conformation selection by NTD; and (v) a degron-tagged Ig2D5 designed to identify the residues of degrons that are recognized and bound by NTD.

Here we show that the presence of NTD is required to degrade damaged proteins and native proteins with degrons, but it does not play an active role in mediating the degradation by LonA of an intrinsically disordered substrate, α-casein. We have also determined the crystal structure of an N-terminal fragment of MtaLonA, and refined it to 2.1 Å resolution. The structure is homologous to the structures of N-terminal fragment of both EcLon and MacLon^19–22^, but not to that of BsLon^*20*^. With the aid of perdeuteration labeling and transverse relaxation optimized NMR spectroscopy (TROSY), we find that the NTD appears to tumble independently of the hexameric core complex, suggesting that the NTD is attached to the hexameric chamber by a flexible linker thus rendering it possible for detailed chemical shift perturbation mapping of substrate binding in the context of full-length MtaLonA. We further structurally characterize the NTD-mediated interactions with unfolded proteins, protein aggregates, or degron-tagged proteins to elucidate its functional role in substrate discrimination. Our results demonstrate that the substrate-binding site of NTD is located at hydrophobic patches of its N-lobe, which recognizes exposed aromatic and hydrophobic residues of damaged proteins and degrons. Using temperature cycling experiments, we investigate how the conformations of substrates are affected in the presence of MtaLonA NTDs, which interact with the initial substrate before subsequent translocation and degradation. Altogether, this work elucidates the substrate recognition mediated by the NTDs of MtaLonA at substantially high resolution and suggests that the flexibly linked NTDs can help survey, discriminate, and selectively capture damaged unfolded protein species or native protein substrates carrying exposed degradation sequence motifs.

## Results

### The NTD is essential for efficient degradation of a thermally unfolded substrate but not intrinsically disordered protein substrate, α-casein

To evaluate the importance of the NTD in the proteolysis activity of MtaLonA, we constructed a MtaLonA variant, AAAP, of which the NTD (residues 1-241) was removed (Fig. 1A), based on the structure of *B. subtilis* LonA. Substrate degradation activity of full-length MtaLonA and AAAP were evaluated using two protein substrates, namely α-casein and Ig2. Native α-casein is an intrinsically disordered substrate of LonA. Ig2 is an all β-stranded protein that becomes partially unfolded at 55 °C (supplementary Fig. 1A) and only then is susceptible to proteolysis by MtaLonA^6^. Full-length MtaLonA degrades both α-casein as well as Ig2^24^. Compared to the full-length MtaLonA, AAAP exhibits a slightly reduced degradation activity against α-casein (Fig. 1B) while its proteolytic activity for Ig2 is severely impaired (Fig. 1C). These results showed that, despite the lack of the NTD, AAAP retains the proteolytic activity for intrinsically disordered protein substrates like α-casein. However, the NTD is essential for MtaLonA to recognize and degrade the thermally damaged substrate, Ig2, indicating the NTD may mediate specific interactions with damaged Ig2 but not α-casein.

**Fig. 1.**
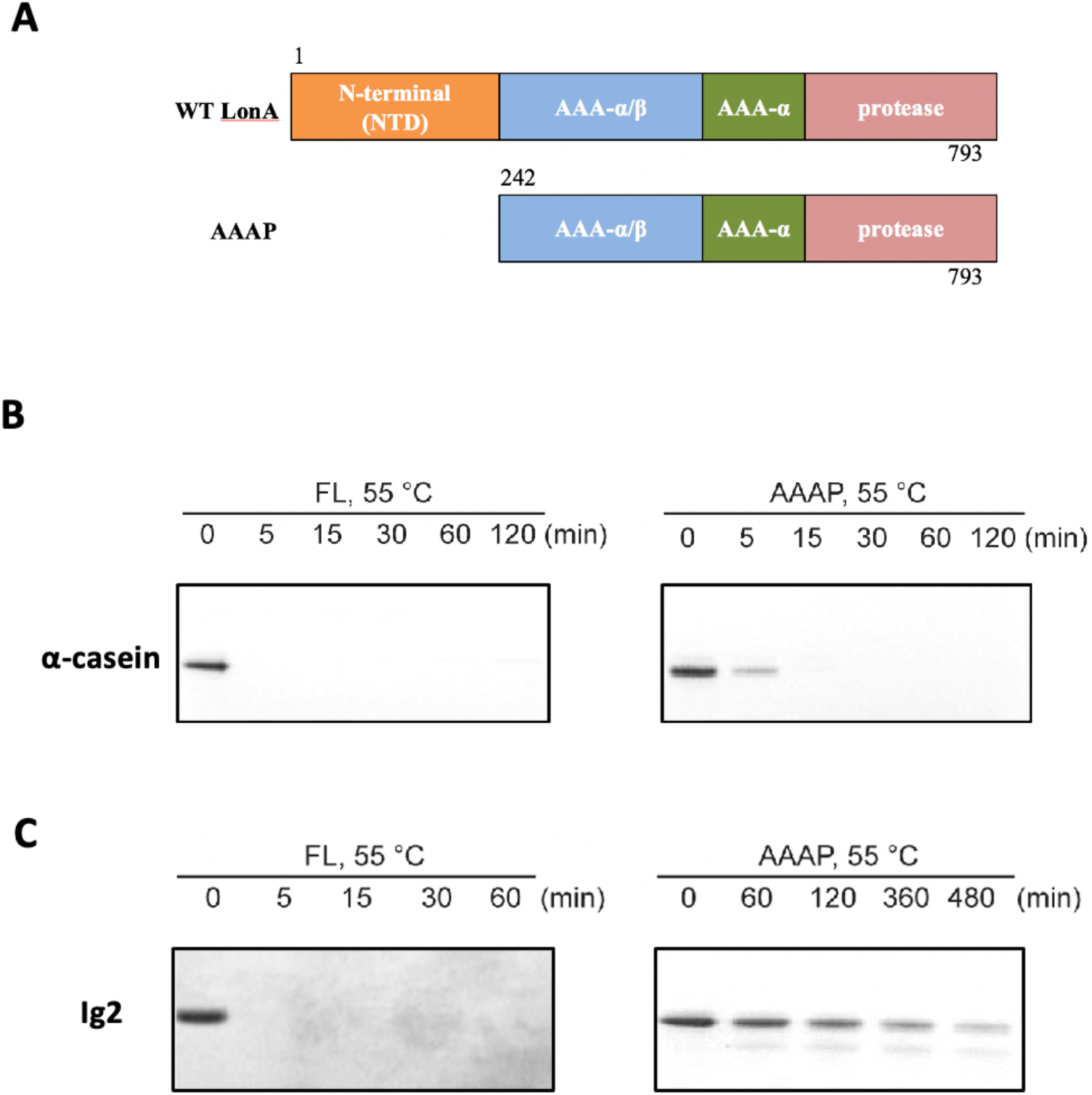
The NTD of Meiothermus taiwanensis LonA (MtaLonA) is essential for efficient degradation of a thermally unfolded substrate, but not the intrinsically disordered substrate, α-casein. (A) Schematic diagrams illustrating domain organization of full-length MtaLonA and the N-terminal domain truncated AAAP. (B) Degradation of α-casein by full-length MtaLonA (FL) and AAAP at 55 °C. (C) Degradation of Ig2 by full-length and AAAP at 55 °C.

### The NTDs tumble independently of the hexameric LonA core in solution

We obtained crystals of the N-terminal fragment NN206 of MtaLonA by *in situ* proteolysis during crystallization (Fig. 2A). The structure was refined to 2.1 Å resolution with a free R factor of 24.9% (crystallographic R factor is 20.9%) (Table 1). NN206 forms a bilobal fold: (i) the N-terminal lobe, which consist of one α-helix and six β-strands, forming three β-sheets (β1/β3/β4/β5, β2/β5/β4 and β1/β6/β5); (ii) the C-terminal lobe, which is is exclusively α-helical, consisting of a five-helix bundle (Figs. 2B). The two lobes are jointed together by an 18-residue linker, which is similar to that of both EcLon and MacLon^19–22^. Based on the result, we also crystallized and solved the structure of the longer N-terminal fragment NN291 (Fig. 2A). The structure of NN291 was refined to 1.7 Å resolution (Table 1), which is similar to NN206 except for a longer C-terminal helix by 4 more residues, i.e., residues 207-210 (Fig. 2C). Despite the lack of spontaneous proteolytic fragmentation in the crystals, no electron density was seen past residue A210 (Fig. 2D).

**Table 1.** Data collection and refinement statistics

|  | NN206 | NN291 |
| --- | --- | --- |
| PDB entry | 7CR9 | 7CRA |
| **Data collection** |  |  |
| Space group | P212121 | P6322 |
| Cell dimensions |  |  |
| *a*, *b*, *c* (Å) | 32.830, 58.296, 198.492 | 86.016, 86.016, 110.126 |
| $\alpha$, $\beta$, $\gamma$ (°) | 90, 90, 90 | 90, 90, 120 |
| Resolution (Å) | 30 - 2.09 (2.16 - 2.09)* | 30 - 1.7 (1.76 - 1.7) |
| *R*<sub>sym</sub> or *R*<sub>merge</sub> | 0.106 (0.937) | 0.069 (0.779) |
| *I*/σ*I* | 17.9 (3.0) | 49.1 (4.7) |
| Completeness (%) | 99.7 (100) | 100 (100) |
| Redundancy | 6.5 (6.6) | 20.9 (21.4) |
| **Refinement** |  |  |
| Resolution (Å) | 29.409 - 2.1 (2.149 - 2.1) | 30 - 1.7 (1.744 - 1.7) |
| *R*<sub>work</sub>/ *R*<sub>free</sub> | 0.209/0.249 | 0.197/0.237 |
| No. atoms |  |  |
| Protein | 3285 | 1684 |
| Ligand (SO<sub>4</sub><sup>2-</sup>) | 0 | 10 |
| Water | 107 | 244 |
| B-factors |  |  |
| Protein | 53.14 | 16.18 |
| Ligand (SO<sub>4</sub><sup>2-</sup>) |  | 27.45 |
| Water | 52.66 | 28.21 |
| R.m.s deviations |  |  |
| Bond lengths (Å) | 0.004 | 0.009 |
| Bond angles (°) | 0.74 | 1.34 |
| Ramachandran |  |  |
| Favored (%) | 96.81 | 96.67 |
| Allowed (%) | 2.46 | 2.38 |
| Outliers (%) | 0.74 | 0.95 |

Number of crystals for each structure should be noted in footnote.
\*Highest resolution shell is shown in parenthesis.

**Fig. 2.**
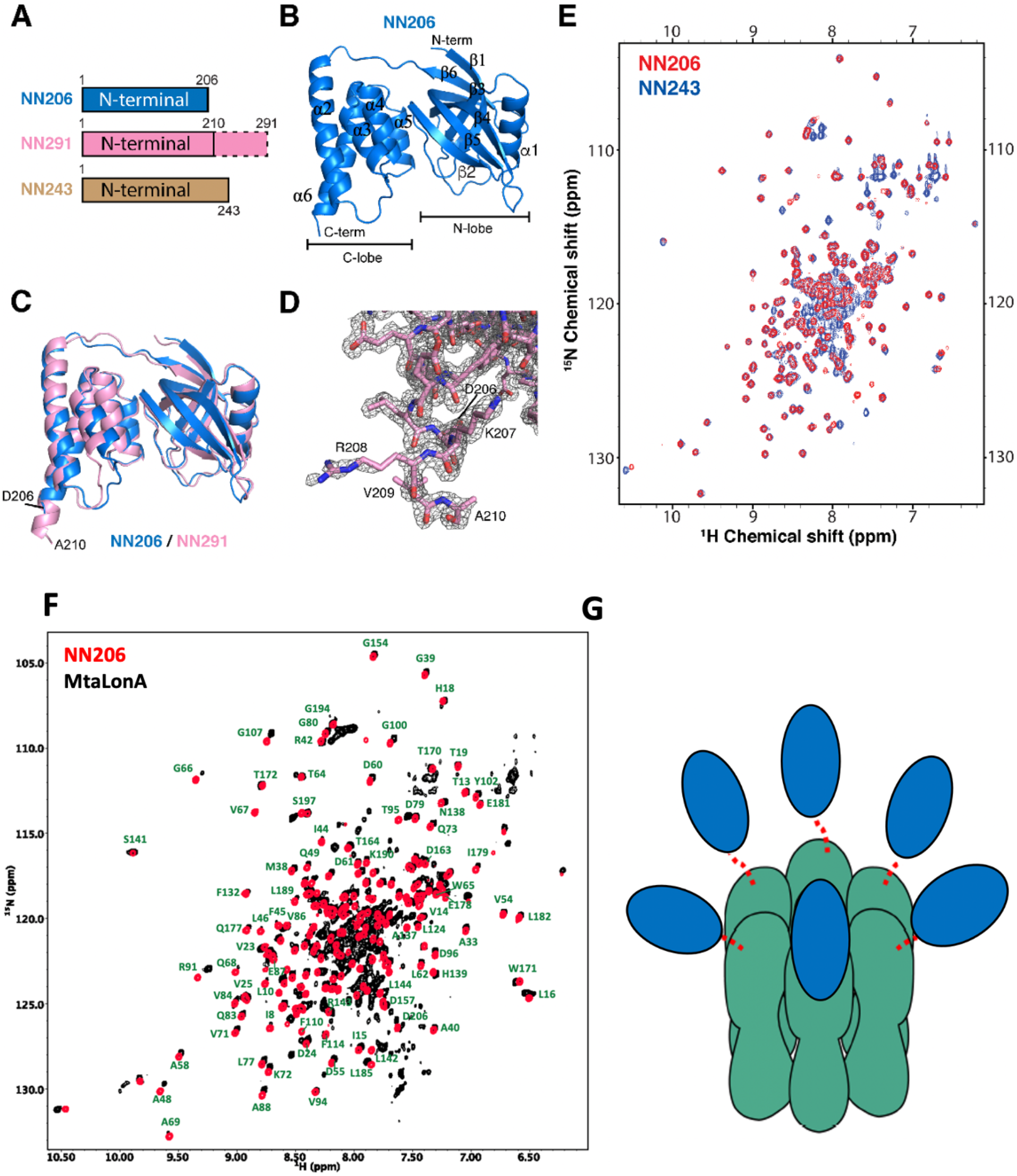
Overall structures of the N-terminal domain of MtaLonA. (A) The three N-terminal constructs NN206, NN291, and NN243. The disordered region is shown in the dashed box. (B) Structure of NN206. (C) Superposition of NN206 (marine) and NN291 (pink) structures. (D) The refined *2F_0_-Fc* electron density map around the C-terminal residue A210 of NN291, contoured at 1.0 *σ*. (E) Overlay of 2D ^1^H–^15^N HSQC NMR spectra of NN206 (red) and NN243 (blue). (F) Comparison of the ^1^H–^15^N TROSY HSQC spectra of NN206 (red) and MtaLonA (black) recorded at 55 °C. The well-dispersed resonances are labeled with residue numbers. About 200 well-resolved correlations are superimposable, suggesting that NTD is loosely linked to the hexameric core via a flexible linker. (G) The six NTD may each connect to the hexameric core of fused AAA+ and protease domains by a flexible ~40-residue linker. The NTDs, and ATPase-Protease chamber are illustrated in blue and green colors, respectively. Red dashed lines represent the flexible linker regions.

Interestingly, the C-terminus of *E. coli* N-terminal fragment forms an extended 40-residue helix^21^. Therefore, we further investigate the structural feature of the corresponding residues 207-243 of MtaLonA using solution-state NMR spectroscopy. Most of the well-dispersed cross-peaks in the two-dimensional ^15^N-^1^H HSQC NMR spectra of NN206 and NN243 can be superimposed, indicating that the well-folded domain structure corresponding to residues 1-206 is not perturbed by the C-terminal extension, residues 207-243 (Fig. 2E). Nonetheless, NN243 exhibits extra intense and poorly dispersed cross-peaks around 7.5-8.5 ppm along the proton dimension that is indicative of a highly disordered peptide segment, which most likely corresponds to the C-terminal extension of NN243. Together, our crystallographic and NMR analyses suggest that the N-terminal domain of MtaLonA forms a bilobal globular structure (residues 1-206) followed by a C-terminal flexible extension (residues 211-243).

LonA forms a large homo-hexameric complex resulting in a very high molecular weight of 0.5 MDa. To verify the dynamic feature of the NTD, we compared the backbone amide ^15^N-^1^H NMR correlations of NN206 and those of full-length MtaLonA at 55 °C. With the aid of perdeuteration labeling and transverse relaxation optimized NMR spectroscopy (TROSY), we could observe over 200 well-resolved ^15^N-^1^H correlations of full-length hexameric MtaLonA at an apparent molecular weight of 0.5 MDa. Importantly, most of these correlations are superimposable to those observed in the NN206 spectrum (Fig. 2F). These results indicate that the NTD adopts the same structure as the isolated NTD while loosely tethered to the near-megadalton hexameric LonA core via a long flexible linker thus leading to an independent, fast tumbling motion to yield highly favorable NMR relaxation properties and thus high-quality multidimensional NMR spectra. We further applied multidimensional heteronuclear NMR experiments to assign the observed cross peaks in the ^1^H–^15^N TROSY HSQC spectrum of NN206 (supplementary Fig. 2). Together, these results suggest that in a LonA complex, the six N-terminal domains may each connect to the hexameric core of fused AAA+ and protease domains by flexible ~40-residue linkers. Therefore, each of the six 24-kDa N-terminal domains may tumble independently of the 350-kDa hexameric core (Fig. 2G).

### The chemical shifts of NTDs are perturbed by thermally unfolded proteins, aggregates, and degron tags

To directly observe discrimination of native or thermally damaged substrate mediated by NTD, we treated NMR samples by a thermal cycle with temperature increasing from 32 °C to 55 °C and then returning to 32 °C. With temperature rising, the exposed hydrophobic areas of thermally unfolded Ig2 is increased, as detected by SYPRO Orange (supplementary Fig. 1A), indicating that the population of Ig2 in unfolded states is increased by thermal denaturation. By contrast, thermal cycling of isolated NTDs exhibited no temperature effect on protein structure monitored by NMR spectroscopy, which may be explained by the high melting temperature (T_m_) of full-length MtaLonA and NN206 (68.0 °C and 85.5 °C, respectively; supplementary Figs. 1A and 1B). Upon addition of a 2-fold molar excess of well-folded Ig2 at 32 °C, no significant chemical shift perturbation (CSP) is observed (Fig. 3A), where only the NTD is isotopically enriched. By increasing temperature from 32 °C to 55 °C for the same NMR sample, a dramatic variation in the 2D spectral features is detected and the resonances undergo a prominent decrease in intensity (Fig. 3B), indicating that the NTD interacts with thermally unfolded Ig2. With temperature dropping from 55 °C to 32 °C, line width of the expected bound-state resonances is too broad to be detected in the presence of binding events (Fig. 3C), suggesting a certain portion of Ig2 still stays in the unfolded or aggregated state. In contrast, the addition of α-casein results in little to no effect on the cross-peaks of residues 1-206 of the NTD (supplementary Fig. 3) thus precluding specific interaction mode between the intrinsically disordered protein α-casein with the NTD of MtaLonA.

**Fig. 3.**
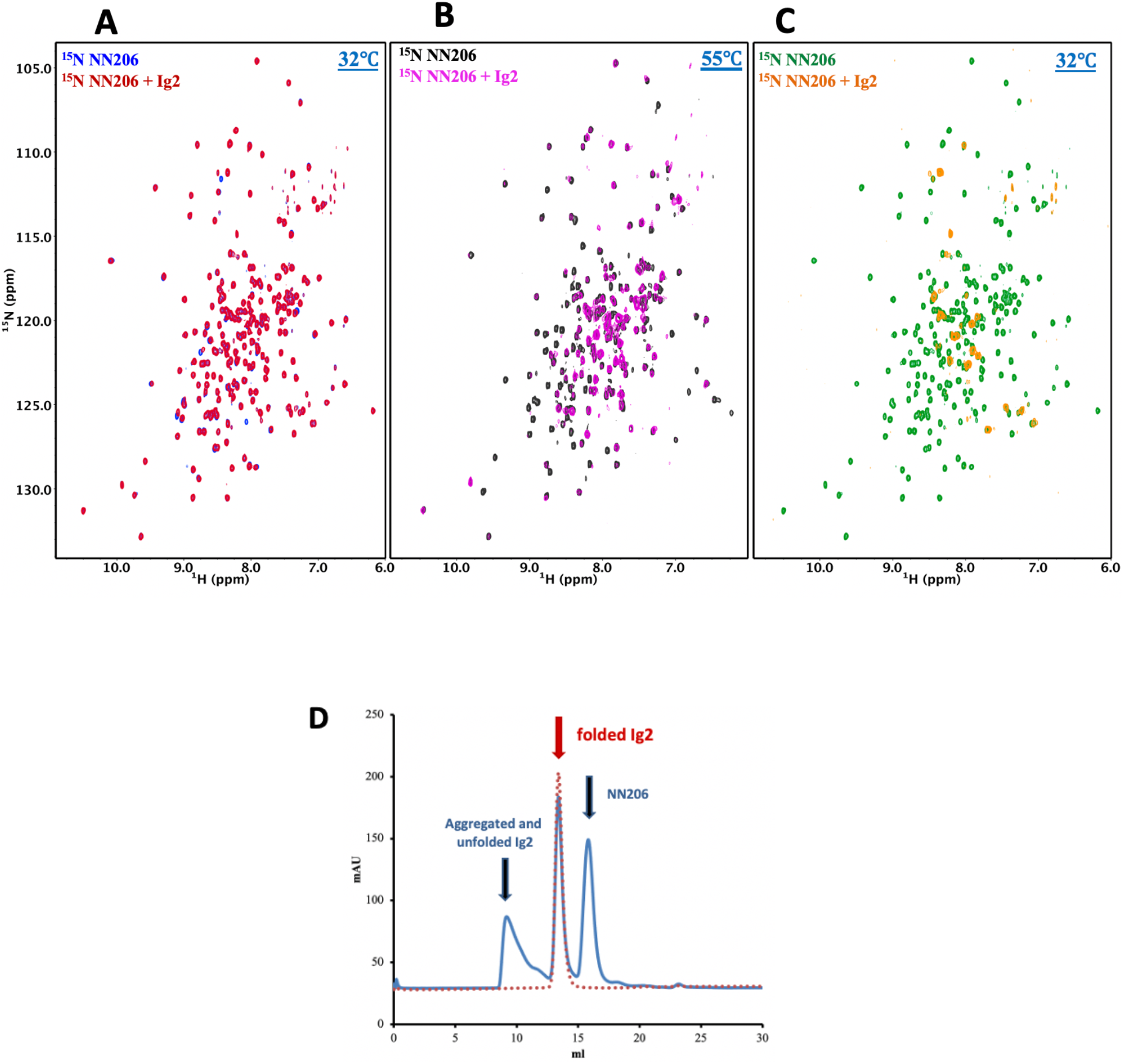
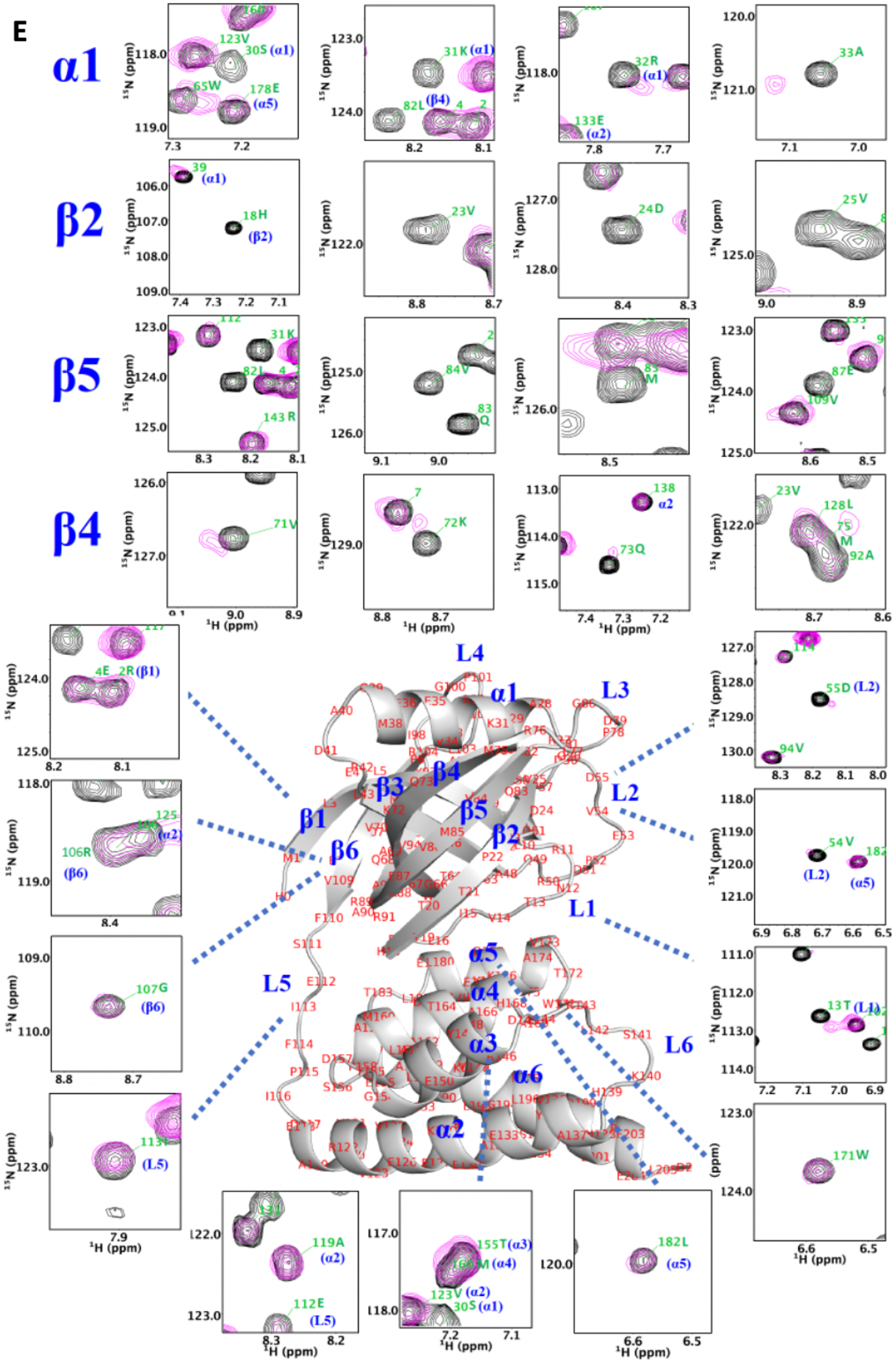
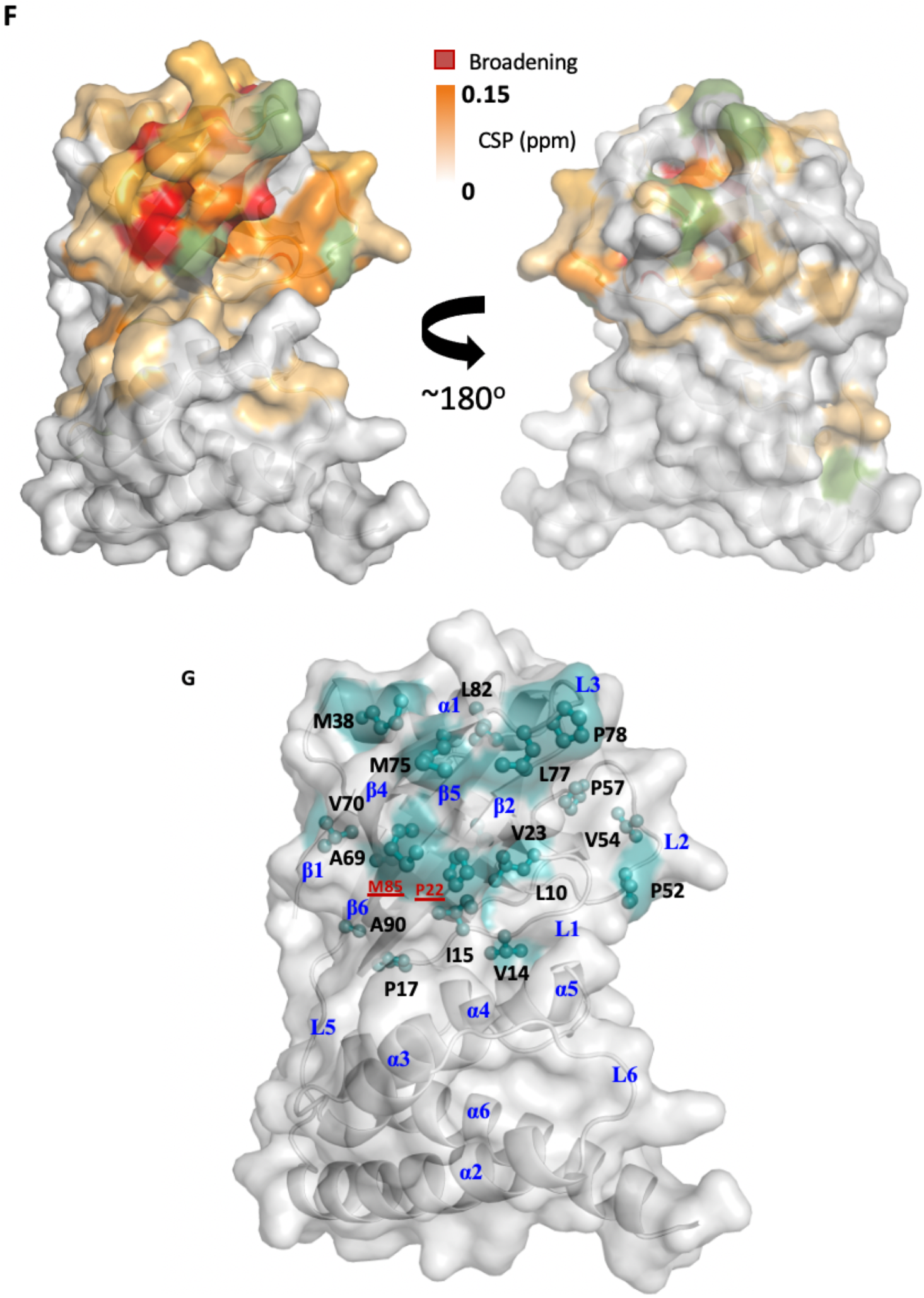
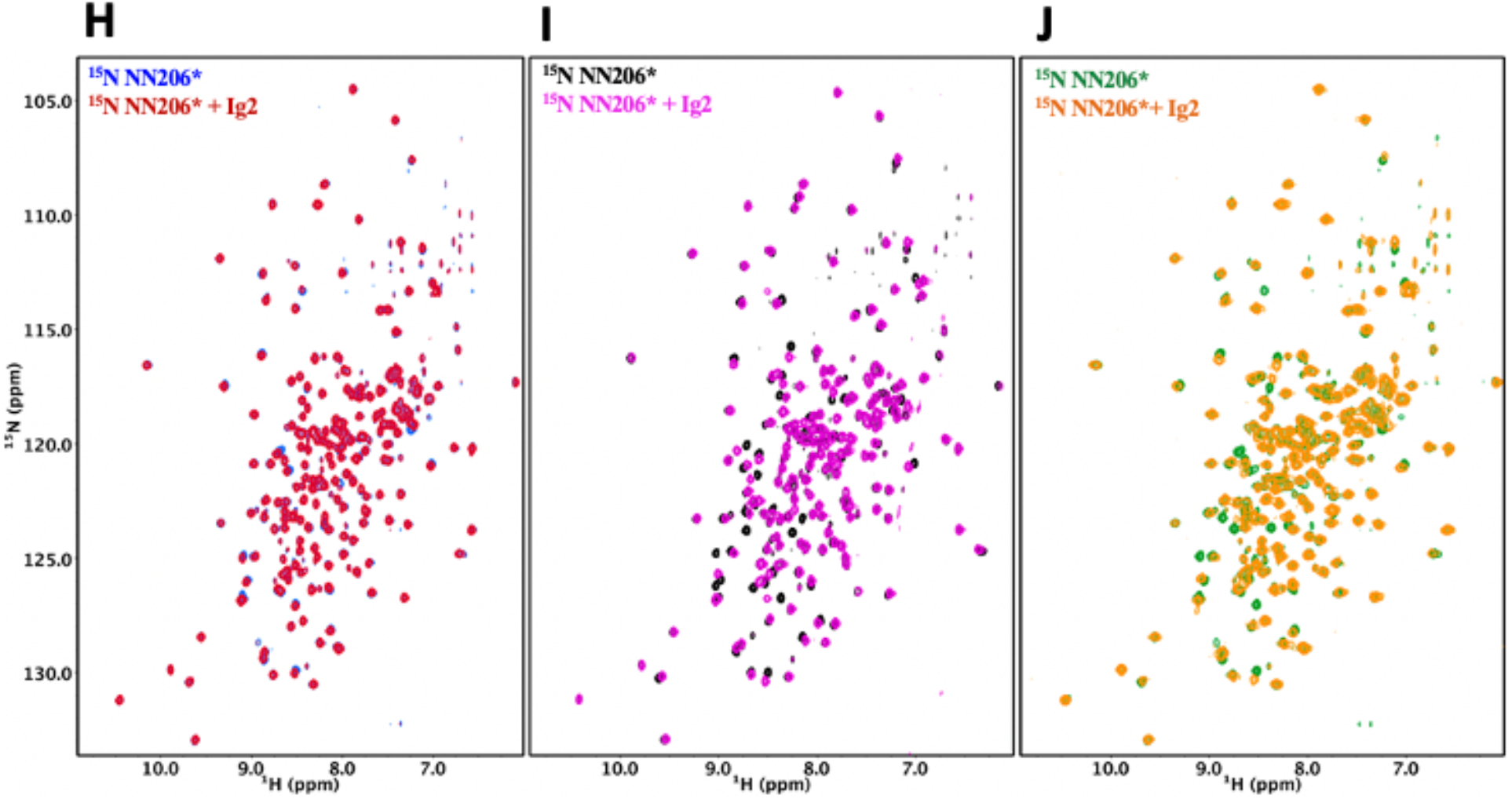
The N-lobe of MtaLonA NTD is involved in recognition of thermally unfolded substrate Ig2. (A) ^1^H–^15^N TROSY HSQC spectra of NTD in the absence (blue) and presence (red) of Ig2 recorded at 32 °C. (B) Comparison of the ^1^H–^15^N TROSY HSQC spectra of NN206 in the apo (black) and Ig2-bound (magenta) states recorded at 55 °C. (C) After one thermal cycle, ^1^H–^15^N TROSY HSQC spectra of NTD in the absence (green) and presence (orange) of Ig2 recorded at 32 °C. (D) Gel filtration profiles of NMR samples treated with thermal cycling shown in blue, and folded Ig2 (red) analyzed by Superdex 200 10/300 GL column. (E) Comparison of the ^1^H–^15^N TROSY HSQC spectra of NN206 in the apo (black) and Ig2-bound (magenta) states. The residues of NTD with the CSPs are part of the interaction interface, located mainly at β‐sheet β2/β5/β4, loop L1, L2, L3 and helix α1. (F) Interaction interface of NTD: Ig2 mapped onto the structure of NTD, based on the spectrum recorded at 55 °C. Proline residue is shown in green. (G) The exposed hydrophobic residues that make up the damaged Ig2-binding sites are shown as balls and sticks. Mutations of two exposed hydrophobic residues P22 and M85, located at the center of β‐sheet β2/β5/β4, lead to reduce the hydrophobicity of the contiguous hydrophobic surface in the N-lobe subdomain. (H) ^1^H–^15^N TROSY HSQC spectra of NTD* (blue) in the absence (blue) and presence (red) of Ig2 recorded through thermal cycling from 32 °C (H) to 55 °C (I) and then returning to 32 °C (J).

To clarify whether overall broadening of the cross-peak signals is due to protein aggregation or precipitation, we performed size-exclusion chromatography (SEC) experiments to examine the NMR sample treated with thermal cycling. The analysis revealed that more than half of Ig2 protein becomes unfolded and aggregated through a thermal cycle while the NTD is still well-folded (Fig. 3D). The NTD-Ig2 aggregate complex showed a dissociable SEC profile, suggesting the NTD-substrate interaction is quite dynamic by nature and the fast substrate dissociation may facilitate the translocation process.

NMR analysis indicated that thermally damaged Ig2 induced significant CSPs and broadened resonances in the NTD N-lobe, which comprise one α-helix and three β-sheets β1/β3/β4/β5, β2/β5/β4 and β1/β6/β5. The residues of NTD with the pronounced CSPs are part of the interaction interface located mainly at helix α1, loop L1, L2, L3 and β-sheet β2/β5/β4 (Figs. 3E and 3F), which consist primarily of hydrophobic residues and are decorated by a number of polar residues (supplementary Fig. 4A). To characterize the binding sites of the NTD interaction with damaged Ig2, we chose to mutate two exposed hydrophobic residues Pro-22 and Met-85 located at the center of β-sheet β2/β5/β4, which form a contiguous hydrophobic surface in the N-lobe subdomain (Fig. 3G and supplementary Fig. 4B). We found that NN206*, containing two specific mutations in the NTD (P22A and M85A; hereafter NN206*), may influence the hydrophobic substrate-binding interaction. The NMR sample containing both NN206* and Ig2 protein was also treated with one thermal cycle. NN206* showed much weak interactions with thermal damaged Ig2 (Figs. 3H-J), confirming that this binding event is mainly contributed by the specific hydrophobic residues with high hydrophobicity in a continuous hydrophobic surface of the N-lobe. Collectively, our results show that the NTD of LonA selectively interacts with protein substrates, via critical bulky hydrophobic residues on the hydrophobic surface of its N-lobe, demonstrating that NTD enables LonA to perform protein quality control by selectively interacting with proteins in damaged or unfolded states.

To further examine how the NTD is involved with protein aggregation, we probed the interactions between the scrambled lysozyme aggregate and MtaLonA NTD by NMR spectroscopy. Native lysozyme is a 14.3-kDa protein (129 amino acids) comprising four disulfide bridges while reduced lysozyme forms loose and flexible aggregates^25^. TCEP (tris (2-carboxyethyl) phosphine; nonthiol-based reducing agent) drives the formation of amorphous lysozyme aggregates observed in this study. Moreover, there is no cysteine residue found in full-length MtaLonA, suggesting TCEP does not affect the structure of NTD. Addition of TCEP-treated denatured lysozymes leaded to substantial spectral changes of NN206 resonances recorded at 32 °C while NN206* showed marginal CSPs marginal CSPs (supplementary Fig. 5A). The structural mapping shows that the affected residues caused by TCEP-treated denatured lysozyme are also located mainly in the N-lobe shown in supplementary Figs. 5B and C. Furthermore, we also examined the degradation activity of LonA against either native or denatured lysozyme, respectively. A gel-based assay showed that full-length MtaLonA can efficiently degrade denatured-reduced lysozyme, but not native lysozyme (supplementary Fig. 5D). Accordingly, AAAP, without the NTD, was inactive against either the scrambled lysozyme aggregate or native lysozyme (supplementary Fig. 5E). Together, these results highlight the key role of NTD, via the N-lobe region, in discriminating and recognizing protein aggregates before subsequent translocation and degradation.

LonA can also degrade natively folded proteins with degradation tags such as sul20 degron^6,26^. Previously, the NTD of *E. coli* LonA was shown to be required for degron binding by chemical cross-linking^7^. To understand how the NTD directly interacts with degrons, a 20-residue peptide (the C-terminal 20 residues of SulA; namely Sul20)^7^ is synthesized and used for NMR titration experiments. Addition of equimolar unlabeled Sul20 causes significant spectral changes of NN206 resonances (Fig. 4A) and the structural mapping shows that the affected residues are again located at β-sheet β2/β5/β4, loop L1, L3 and helix α1 of the N-lobe subdomain (Fig. 4B). Mutations of P22 and M85 located at the center of the hydrophobic surface to moderately hydrophobic Ala lead to much weak interactions of NN206* with Sul20 peptide (supplementary Fig. 6A), suggesting that specific hydrophobic residues of beta-stranded β2/β5/β4 located at the N-lobe of MtaLon NTD play a critical role on degrons recognition.

**Fig. 4.**
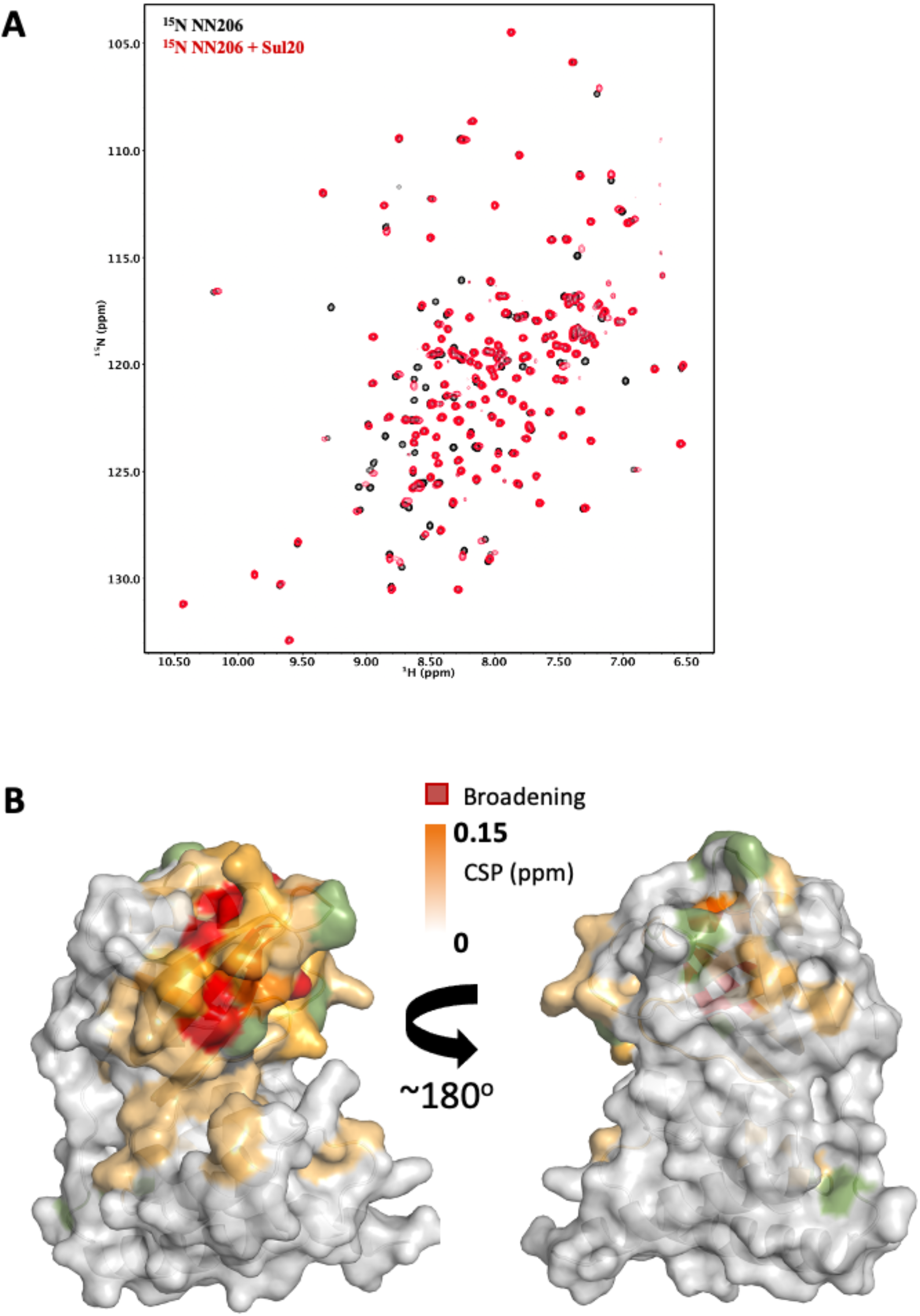
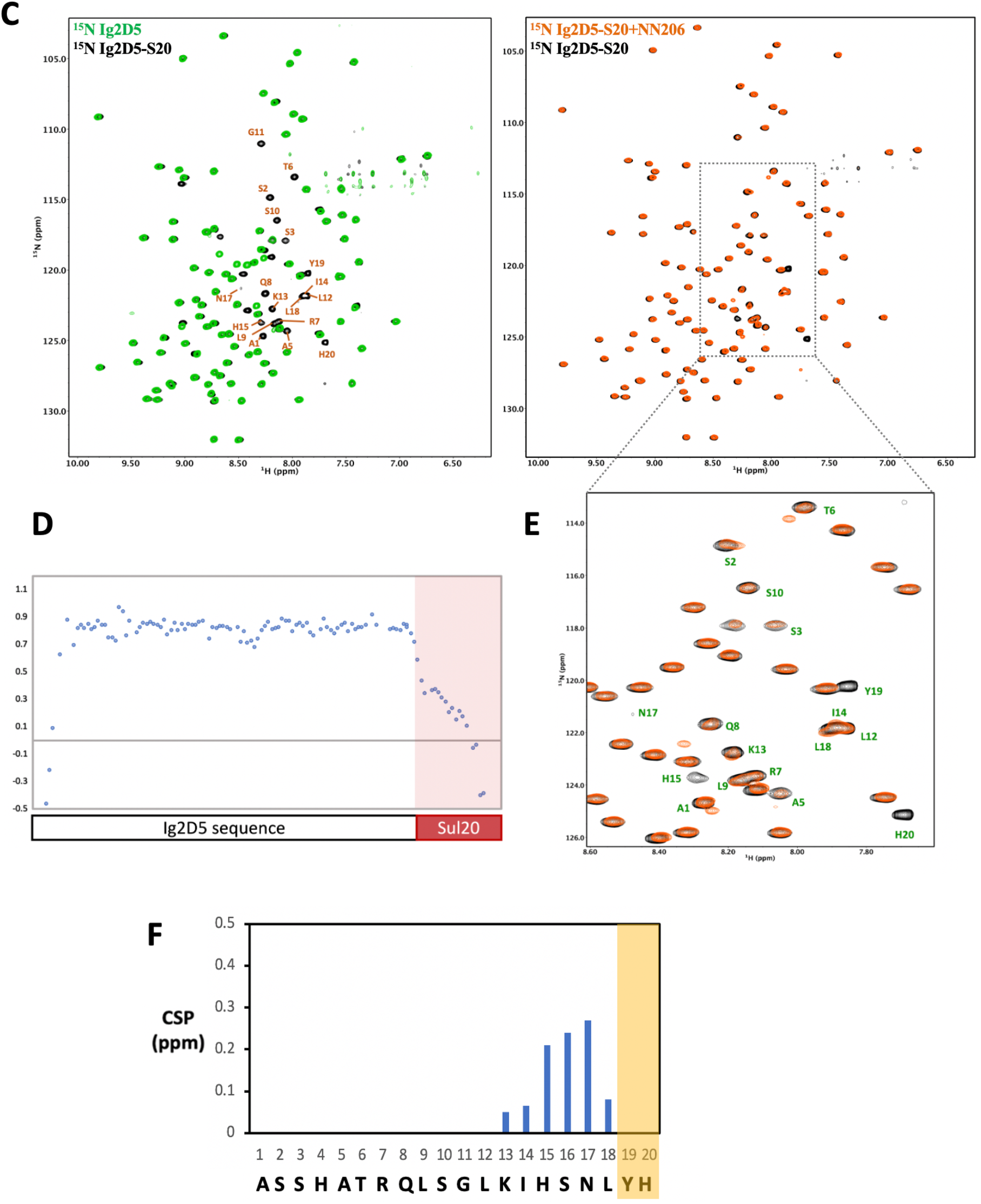
Interaction of MtaLonA NTDs with degron tag Sul20. (A) Overlay of the ^1^H–^15^N TROSY HSQC spectra of NN206 in the absence (black) and presence (red) of Sul20 peptide. (B) Structural mapping of the chemical shift perturbations (orange) of NN206 caused by Sul20 peptide. (C) Overlay of the ^1^H–^15^N TROSY HSQC spectra of Ig2D5 (green) and Ig2D5-S20 (black); overlay of the ^1^H–^15^N TROSY HSQC spectra of Ig2D5-S20 in the absence (black) and in the presence (orange) of an equimolar amount of NN206. (D) [1H]–15N nuclear Overhauser effect (NOE) values of Ig2D5-S20. (E) Expanded region of the ^1^H–^15^N TROSY HSQC spectra of Ig2D5-S20 in the absence (black) and in the presence (orange) of NN206. (F) CSPs of amide moieties of 50 μM Ig2D5-S20 after binding 50 μM unlabeled NN206, plotted against Sul20 residue number. Residues Y19 and H20 are strongly affected by the binding event based on the resonance broadening beyond detection.

To characterize the effect of MtaLon NTD on degrons further, we appended Sul20 to the C terminus of a protein substrate Ig2D5 (domains 5 of the gelation factor from *Dictyostelium discoideum*, hereafter abbreviated as Ig2D5) and investigated the conformational changes experienced by Sul20 upon binding the NTD. The NMR spectra of Ig2D5 and Ig2D5-S20 can be well superimposed, indicating that the well-folded domain structure corresponding to Ig2D5 is not perturbed by the C-terminal Sul20 (Fig. 4C). Furthermore, Ig2D5-S20 exhibits additional cross-peaks that well correspond to Sul20. Elevated [^1^H]–^15^N nuclear Overhauser effect (NOE) values for most residues are from Ig2D5 (Fig. 4D), along with the large increase in chemical shift dispersion. Lower [^1^H]–^15^N NOEs for Sul20 residues are due to the increased flexibility on ps-ns timescale. By addition of the unlabeled NN206 to isotopically enriched Ig2D5-S20, we observed chemical shift changes and broadened resonances of C-terminal Sul20 (Figs. 4E and 4F), suggesting that residues Y19 and H20 are strongly affected by the binding event based on the resonance broadening beyond detection. The NMR spectrum of Ig2D5-S20 in the presence of an equimolar amount of unlabeled NN206* shows a much less pronounced chemical shift effect (supplementary Fig. 6B) than wild-type NN206 binding to Ig2D5-S20, indicating that Sul20 tag can be recognized by the hydrophobic patches of N-lobe of MtaLon NTD.

### The thermally unfolded conformations of substrate proteins are stabilized by engagement with NTDs

To understand further selective recognition of protein substrate in the unfolded states by the NTD, we ask whether the unfolding-refolding process of protein may be modulated by interacting with the NTD. To investigate how the conformations of substrates are influenced in the presence of NTDs, we constructed a single-domain protein substrate Ig2D5 (domains 5 of the gelation factor from *Dictyostelium discoideum*) for NMR experiments due to its high-quality NMR spectrum (supplementary Fig. 7A). About 96% of the backbone resonances of Ig2D5 can be assigned by multidimensional heteronuclear NMR experiments. Ig2D5 is a β-sheet protein with Ig-like fold and the T_m_ of substrate Ig2D5 is similar to Ig2 (supplementary Figs. 7B and 7C). Here we applied NMR spectroscopy to investigate unfolding-refolding equilibria of the substrate Ig2D5 induced by temperature cycling while MtaLon NTD remains stable through the entire temperature cycling. The NMR samples containing isotopically enriched Ig2D5 in the absence and presence of unlabeled NN206 were heated from 32 °C to 60 °C and then cooled to 32 °C. Based on the intensity changes of resonances corresponding to native Ig2D5, we estimated that isolated Ig2D5 is largely unfolded at 60 °C (Figs. 5A and B; the spectra are shown in green) and ~72% becomes folded after a thermal cycling (Fig. 5C). However, in the presence of equimolar NN206, less than 15% Ig2D5 is folded after one thermal cycle (Figs. 5C; the spectrum is shown in blue). Similar results were obtained using equimolar full-length MtaLonA catalytic mutant S678A (LonA*) (Figs. 5C; the spectrum is shown in red). However, addition of equimolar AAAP* (residues 242-793 with catalytic mutant S678A), Ig2D5 is largely unfolded at 60 °C (Fig. 5B; the spectrum is shown in black) and ~75% becomes folded after thermal cycling (Fig. 5C; the spectrum is in black). The results reveal that the unfolded state of Ig2D5 is engaged with MtaLon NTD and the temperature-induced unfolding-refolding of substrate is significantly affected in the presence of MtaLon NTDs (Fig. 5D), potentially priming it for subsequent translocation process. Accordingly, we also examined how the unfolding-refolding of Ig2D5 can be affected by NN206*. After one thermal cycle, about 75% of Ig2D5 becomes folded in the presence of equimolar NN206* (Fig. 5E and supplementary Figs. 7D, E and F). Upon addition of eightfold excess of NN206*, about 45% of Ig2D5 becomes folded, suggesting that NN206* interacts with denatured protein substrate weakly. These results demonstrate that the conformations of substrate are affected by the hydrophobic substrate-binding interaction and the thermally unfolded states of substrate proteins can be stabilized by engagement with the NTDs. Thus, by selectively interacting with hydrophobic residues exposed in the thermally unfolded states of substrate proteins NTDs prominently perturb temperature-induced unfolding-refolding process of substrate.

**Fig. 5.**
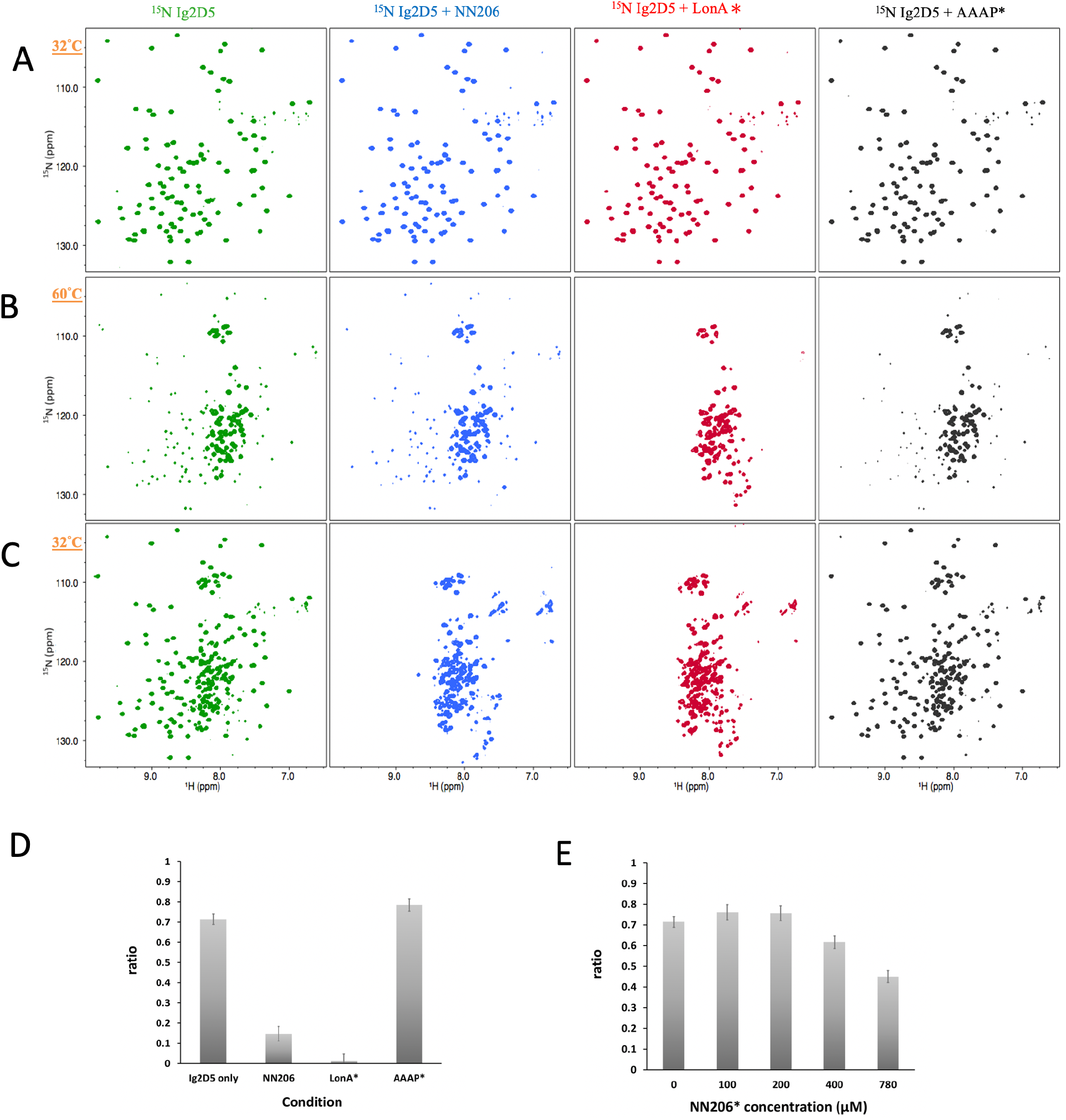
The thermodynamics of Ig2D5 unfolding-refolding is affected in the presence of MtaLonA NTDs. 2D ^1^H–^15^N TROSY HSQC NMR spectra of apo Ig2D5(green), Ig2D5 titrated with NN206(blue), LonA (red), or AAAP (black) are recorded with a thermal cycle starting from 32 °C (A) to 60 °C (B), and then returning to 32 °C (C). (D) Folded ratio of apo Ig2D5, Ig2D5 titrated with NN206(blue), LonA (red), or AAAP (black) after a thermal cycle. (E) Folded ratio of apo Ig2D5, Ig2D5 titrated with different concentrations of NN206* after one thermal cycle.

## Discussion

In this work we used α-casein and Ig2, which represent two types of substrates. Native α-casein is an intrinsically disordered protein; by contrast, the native Ig2 is a β-sheet protein with T_m_ at 46 °C (Supplementary Fig. 1). We show that the construct AAAP devoid of the NTD (residues 242-793, ATPase and protease domains), which retains wild-type ATPase and peptidase activities in LonA, lacks the degradation activity against thermally denatured Ig2 while exhibits a wild-type-like activity against α-casein (Fig.1). Therefore, the NTD is specific for denatured or damaged protein substrates, whose hydrophobic core regions normally buried in the folded state become exposed, but not for intrinsically disordered substrates like casein, which contains mainly charged or polar residues and lack hydrophobic regions in the protein sequence. Indeed, it has been shown that LonA prefers substrates rich in hydrophobic residues^26^.

Our crystallographic results of NN206 and NN291 are consistent with previous findings that the ~40-residue region between the NTD and ATPase-Protease domains is structurally flexible with multiple accessible protease sites^9,20,27,28^. This notion is also corroborated by our high-resolution NMR analyses on the NTD both in isolation and as a part of the full-length protein, demonstrating the independent, fast tumbling motion of the NTDs in the ~0.5 MDa hexameric assembly of LonA. Moreover, their highly favorable NMR relaxation properties and resulting high-quality multidimensional NMR spectra have allowed us to analyze the interaction of the NTD with unfolded proteins, protein aggregates, degron tags, and intrinsically disordered substrates. Our work provides atomic details of the NTD-mediated substrate discrimination by temperature-based switching of the native folded protein to its unfolded state in an NMR sample. NMR characterization indicates NTD does not interact with well-folded Ig2, native lysozyme or intrinsically disordered α-casein, whereas unfolded proteins, aggregates and degron tags can elicit pronounced chemical shift changes to the N-lobe of the NTD, which is subsequently confirmed by structure-based mutagenesis. Collectively, these results suggests that LonA has two substrate-interacting modes: (1) the NTD-non-requiring mode for intrinsically disordered substrates lacking hydrophobic-rich region, which may be engaged directly with the pore-loops in the LonA assembly without involving the six NTDs; (2) the NTD-requiring mode for recognizing and trapping damaged/denatured protein substrates with exposed hydrophobic regions before their engagement with the pore-loops of the LonA assembly. The key role of NTD is to enable LonA to perform protein quality control to selectively capture and degrade proteins in damaged unfolded states.

By analyzing the NTD of MtaLonA interactions with damaged Ig2, the scrambled lysozyme and Sul 20 peptide, we conclude that these binding events are mediated by two hydrophobic patches, which comprise (i) Leu-10, Ile-15, Pro-22, Val-23, and Met-85 (termed hpI in the following); (ii) Met-75, Leu-77, Pro-78 and Leu-82 (termed hpII) (Fig. 6A). Our results also demonstrate that a double mutant NN206* (Pro-22 and Met-85 replaced by Ala), resulting in decreased hydrophobicity of hpI, has a significant effect on the ability of MtaLonA NTD to bind to damaged or misfolded proteins, indicating that the bulky hydrophobic residues on hydrophobic patches are essential determinants for substrate recognition and discrimination. We also examine the exposed hydrophobic residues of β-sheet β2/β5/β4, loop L1 and loop L3 located at the N-lobes of other reported structures of LonA NTDs^19–22^. The N-terminal fragment from *B. subtilis* LonA was previously reported to adopt a domain-swapped dimer where the N-lobe of one monomer is positioned next to the C-lobe of the other monomer^20^. In contrast, the structures of the N-terminal fragment from MtaLon, EcLon and MacLon showed that the N- and the C-lobes are joined together via a short linker (supplementary Fig. 8A and B). Based on structural alignments (supplementary Fig. 8C), the N-lobe of EcLon has almost the same pattern of exposed hydrophobic residues as that of MtaLon NTD (Fig. 6B). Interestingly, The NTD of MacLon shows merged hp I and II while the hpI of BsLon NTD is a little distant from hpII (Figs. 6C and D). The structural comparison reveals that, similarly to MtaLon NTD, all three reported structures exhibit exposed hp I and II with slight variations in their shapes and sizes, suggesting that the conserved hydrophobic patches at the N-lobes may be potentially responsible for conferring substrate selectivity toward degrons, misfolded and damaged proteins.

**Fig. 6.**
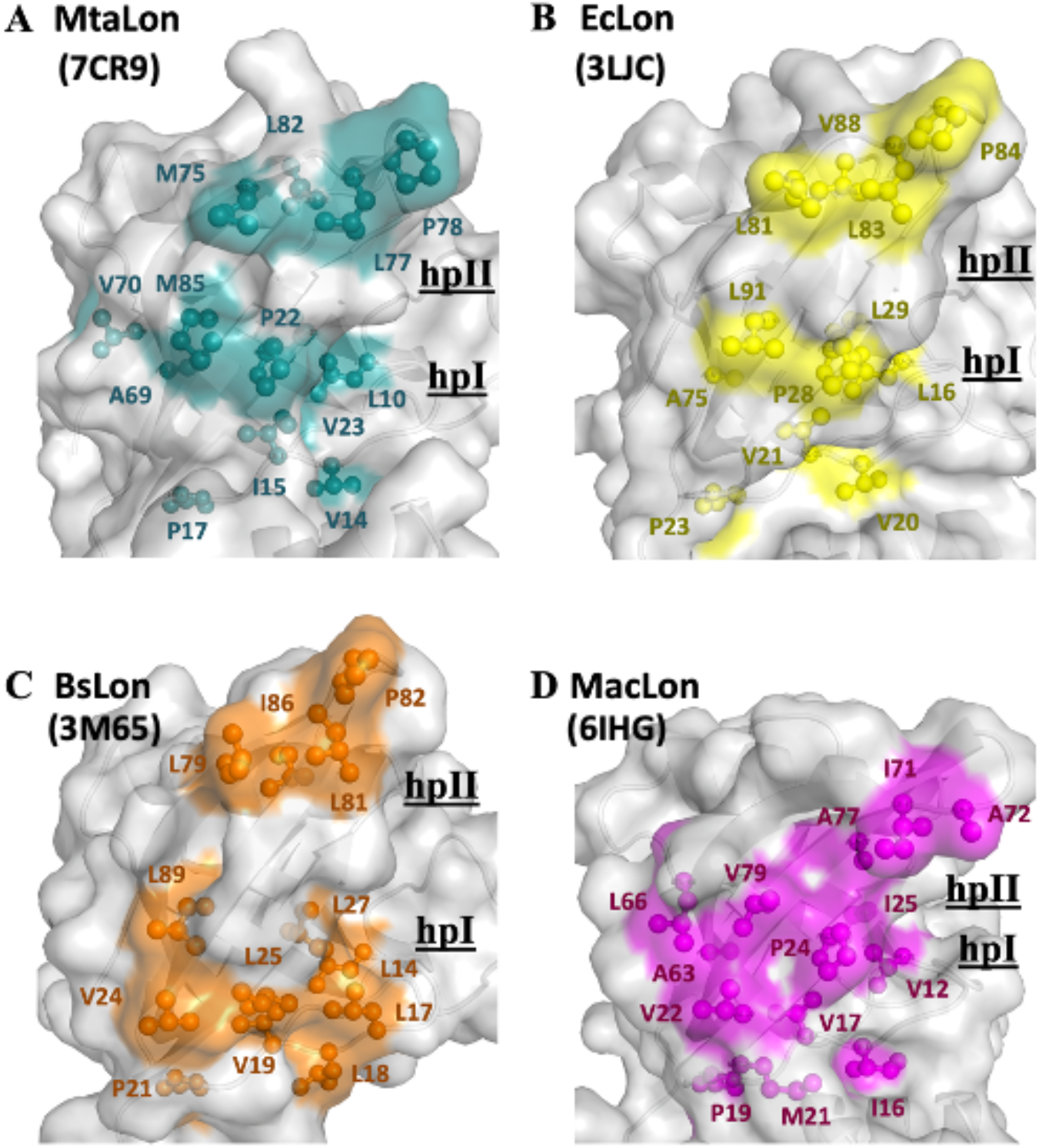
The NTD of MtaLonA selectively interacts with unfolded proteins, protein aggregates, and degron-tagged proteins via two hydrophobic patches of its N-lobe. (A) Two hydrophobic patches located at the N-lobe of MtaLonA: (i) Leu-10, Ile-15, Pro-22, Val-23, and Met-85 (termed hpI in the following); (ii) Met-75, Leu-77, Pro-78 and Leu-82 (termed hpII). (B) Hp I and II of EcLonA: (i) Leu-16, Val-21, Pro-28, Leu-29, and Leu-91; (ii) Leu-81, Leu-83, Pro-84 and Val-88. (C) Hp I and II of BsLonA: (i) Leu-14, Leu-17, Leu-18, Val-19, Val-24, Leu-25, Leu-27 and Leu-89; (ii) Leu-79, Leu-81, Pro-82 and Ile-86. (D) merged Hp I and II of MacLonA: (i) Val-12, Val-17, Val-22, Pro-24, Ile-25, Leu-66 and Val-79; (ii) Ile-71, Ala-72 and Ala-77. The conserved hydrophobic patches at the N-lobes may be potentially responsible for conferring substrate selectivity toward degrons, misfolded and damaged proteins. The PDB codes are indicated in parentheses.

It has previously been known that LonA can recognize a degron-tagged protein by binding to its largely exposed hydrophobic residues. Here, our results show the C-terminal Sul20 of Ig2D5-S20 has substantially low heteronuclear NOE values, reflecting this region undergoes rapid local motion and is solvent-exposed in solution. Therefore, NTD of MtaLonA can easily recognize and interact with the C-terminal aromatic and hydrophobic residues of Sul20. This finding indicates that a pair of aromatic residues consisting of Y19 and H20 in Sul20 play crucial roles in the hydrophobic interactions with residues P22 and M85 of LonA NTD, which is verified with a double mutant NN206*. Residues Pro and Met exhibit higher values of hydrophobicity than moderately hydrophobic Ala and mutations of both P22 and M85, located at the center of the contiguous hydrophobic surface, result in much weak interactions of NN206* with damaged Ig2, the scrambled lysozyme and Sul20 peptide, highlighting the importance of direct interaction with exposed and contiguous surface with high hydrophobicity in the recognition of protein substrates by the NTD of MtaLonA. Furthermore, by selectively binding to hydrophobic residues exposed in the thermally unfolded states of proteins, the NTDs prominently stabilize substrate’s unfolded conformation. These results show that the NTD-substrates interactions involve the hydrophobic residues on both partners, suggesting aromatic and hydrophobic side chains, which are usually buried in the native proteins, are important recognition determinants for the NTD to enable MtaLonA to selectively capture and degrade proteins in a damaged unfolded state or with degron tags.

Our results directly demonstrate that thermally damaged proteins and degrons induce chemical shift changes and broadened resonances located mainly in the two hydrophobic patches of N-terminal lobe but we do not find evidence that the C-lobe of the NTD is involved in substrate interaction. However, addition of thermally unfolded Ig2 into U-^2^H/^15^N-labelled full-length LonA*, affected residues are shown in both N- and C-lobes of LonA* NTD at 55 °C (Figs. 7A and B). Compared ^1^H–^15^N TROSY HSQC spectra of NN206 and MtaLonA* in the presence of thermally unfolded protein Ig2, the signals from the C-lobe of MtaLonA* NTD are missing due to line broadening, suggesting others NTDs of LonA* hexamer may join to interact with substrates. All together, these results reveal the free molecular tumbling of the NTD and its substrate binding site, based on which a model involving multi-NTD interactions, may be proposed for substrate recognition. The flexibly linked NTD of the hexametric LonA is swaying from side to side and back and forth to survey, recognize, and trap substrates with exposed hydrophobic sequences (Fig. 7C).

**Fig. 7.**
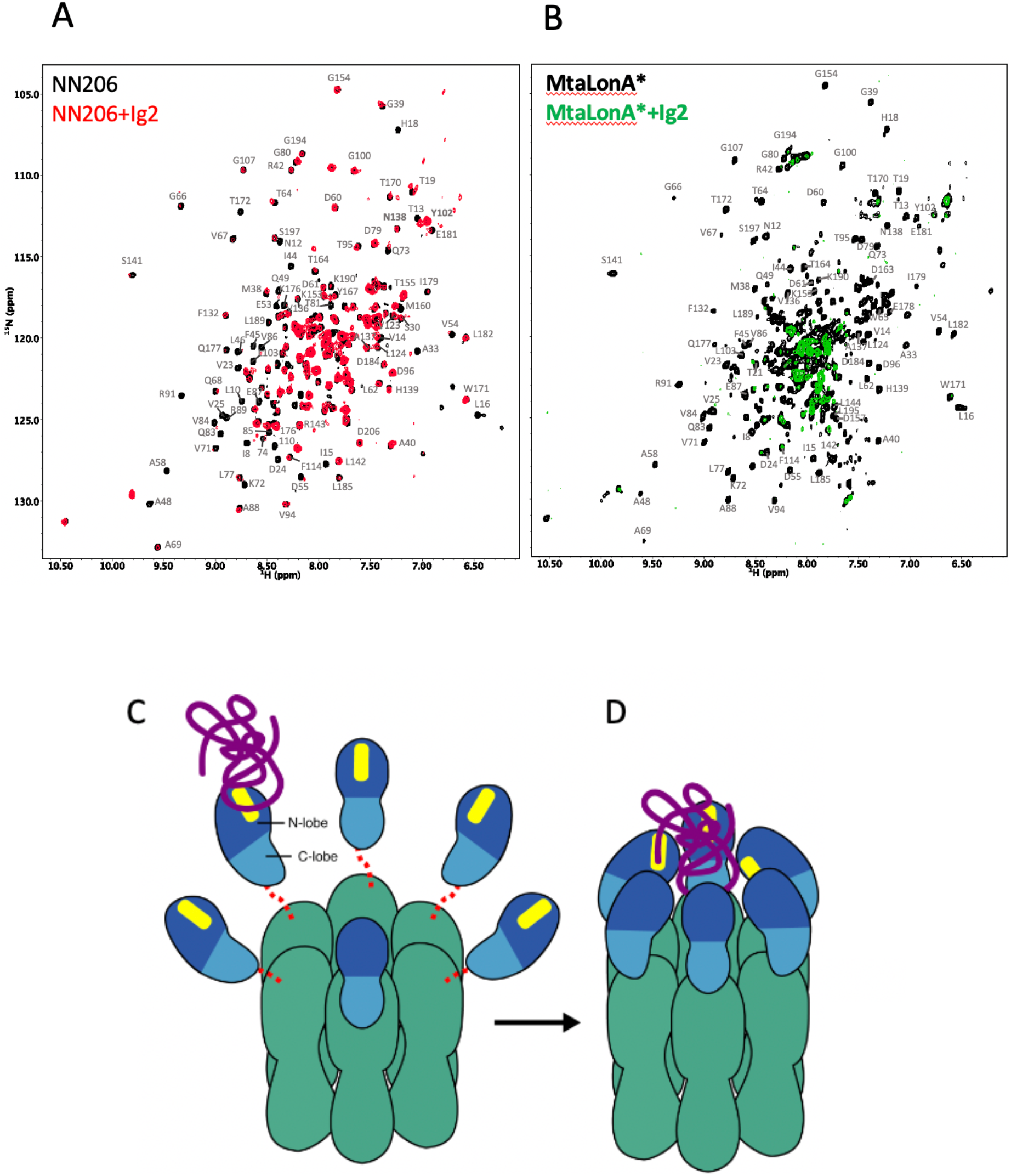
Proposed model for NTD-mediated substrate interaction. (A) ^1^H–^15^N TROSY HSQC spectra of NTD in the absence (black) and presence (red) of unlabeled Ig2 recorded at 55 °C. (B) ^1^H–^15^N TROSY HSQC spectra of LonA* in the absence (black) and presence (green) of unlabeled Ig2 recorded at 55 °C. (C) The flexibly linked NTD of the hexametric LonA is swaying from side to side and back and forth to survey, recognize, and trap substrates with exposed hydrophobic sequences. Substrate, N-lobe, Clobe of NTDs, and the ATPase-Protease chamber are illustrated in purple, marine, blue and green colors, respectively. The substrate-binding patches are depicted in yellow. Red dashed lines represent the flexible linker regions. (D) After initial binding, other NTDs may join to increase the avidity of the LonA-substrate interaction. The substrate polypeptide chain is pulled inside the chamber and undergoes proteolysis by protease modules of LonA.

After initial binding, other NTDs may join to increase the avidity of the LonA-substrate interaction (Fig. 7D); the engagement of a substrate protein by multiple NTDs also serves to facilitate effective substrate unfolding and translocation mediated by the coordinated movements of the pore-loops in the ATPase modules of the LonA chamber, powered by cycles of ATP binding and hydrolysis. Finally, the substrate polypeptide chain is pulled inside the chamber and undergoes proteolysis by protease modules of LonA.

## Materials and Methods

### Peptide

Sul20 peptide (Sequence: ASSHATRQLSGLKIHSNLYH) was synthesized by GenScript (https://www.genscript.com/) at >95% purity.

### Cloning, protein expression and purification

The plasmids expressing for full-length MtaLonA (1-793 residues) or AAAP (242-793 residues) with a C-terminal 6xHis-tag were transformed into *E.coli* BL21(DE3) cells^18^. Site-directed mutagenesis was performed using the Quickchange kit (Stratagene). NN206 (1-206 residues), NN243 (1-243 residues) and NN291 (1-291 residues) were cloned into pET-modified vector with a tobacco etch virus (TEV) cleavage site. The three resultant plasmids, encoding the proteins NN206, NN243 and NN291 with the N-terminal His-tag, were transformed into *E.coli* BL21(DE3) cells. The target proteins synthesis was induced with 0.5 mM isopropyl-thio-β-D-galactoside (IPTG) at an absorbance at 600 nm (OD_600_) ~0.6 at 28 °C. The target proteins were purified with nickel chelate affinity resins (Ni-NTA, Qiagen). The protein samples were collected and purified by a Superdex 200 (GE Healthcare) column^17^. MtaLonA and AAAP proteins were collected and further purified on a MonoQ (GE Healthcare) chromatography. For NN206, NN243 and NN291 purification, the N-terminal His-tagged proteins were cleaved by TEV protease for overnight at 4 °C and then reloaded onto Ni-NTA to remove TEV protease. The flow-through fraction containing target proteins were collected and purified with Superdex 75 (GE Healthcare) chromatography. Ig2 (domains 5 and 6 of the gelation factor ABP-120 of *Dictyostelium discoideum*)^29^ and Ig2D5-S20 (Ig2 fused with Sul20) were cloned into pET28a(+)tev for generation of pET28a(+)tev-Ig2 and pET28a(+)tev-Ig2D5-20. The recombinant protein was induced by 0.5 mM IPTG at OD_600_ ~0.6 for 16 hours. The purifications of Ig2, Ig2D5 and Ig2D5-20 is the same as NN206, NN243 and NN291. Isotopically labeled samples for NMR studies were prepared by growing the cells in minimal (M9) medium. The full-length MtaLonA were prepared by supplementing the growing medium with 1 g l^−1^ of ^15^NH4Cl and 4 g l^−1^ of ^2^H_7_/^12^C_6_-glucose in 99.8%-^2^H_2_O (Sigma-Aldrich).

### Substrate degradation assays

α-casein (Sigma, USA) and Ig2 were used as the substrates in these assays^17^. 4 μM substrate proteins were incubated with 0.4 μM MtaLonA (hexamer) or AAAP or mutations at 55 °C. The degradation reactions were halted by protein sample dye and heat inactivation at 95 °C for 5 minutes. The treated reaction mixtures were then analyzed by SDS-PAGE.

### Protein crystallization

Crystallizations of NN206 and NN291 were performed at 295K by hanging drop vapor-diffusion method. For Crystallization of NN206, in situ proteolysis was performed^30^. 1 μl of MtaLonA (10 mg/ml) plus trypsin in ratio 1000:1 was mixed with 1 μl of 0.1 M Tris-HCl (pH 7.5), 0.2 M Sodium acetate and 30% PEG 4000. The NN206 crystals appeared in the mixed drop after 3 days. The crystals of NN291 were grown by mixing with 1 μl of NN291 protein sample (15 mg/mL) and 1 μl of well solution containing 0.1 M MES (pH 5.8) and 0.8 M ammonium sulfate. Crystals grew to 0.2-0.3 mm over two weeks. Crystals of NN206 and NN291 were cryoprotected with 20% glycerol or 20% ethylene glycol before data collection.

### Structure determination

The data sets of NN206 and NN291 were collected at the beamlines BL-13C1 of National Synchrotron Radiation Research Center (Taiwan) and BL-1A of Photon Factory (Japan), respectively. All Data sets were processed by HKL2000^31^. The structure of NN206 was solved by molecular replacement with the program Phaser^32^ using the structures of the N-terminal fragment of *B. subtilis* LonA (residues 4-116, PDB code 3M65) and the N-terminal fragment of *E. coli* LonA (residues 120-210, PDB code 3LJC) as the search models. The structure of NN291 was determined by molecular replacement using NN206 structure. All initial models were automatically rebuilt using the program AutoBuild^33^. Subsequently, these structures were refined by manual refitting in Coot^34^ and performance of refinement using the program Refmac5^35^. Crystallographic and refinement statistics are listed in Table 1. All structure figures were made using PyMOL (Version 1.3, Schrödinger, LLC). The atomic coordinates and structure factors for NN206 and NN291 have been deposited in the Protein Data Bank (http://www.rcsb.org/) with the accession numbers 7CR9 and 7CRA, respectively.

### Thermal shift assay

Thermal shift assay was carried out in qPCR 8-strip tubes (Gunster Biotech) using SYBRO orange (Life Technologies) as dye. For each reaction, 5 μM purified MtaLonA, NN206 or Ig2 was mixed with assay buffer (50 mM NaPi, 2 mM β-Me, pH 6.5) and SYPRO Orange dye (5X final concentration) in 45 μL total volume/well. 100 μM Ig2D5 was mixed with assay buffer and SYPRO Orange dye (10X final concentration) in 45 μL total volume/well. Samples were heated at a rate of 0.5 °C per minute from 25 °C to 95 °C and fluorescence signals were recorded with the CFX Connect Real-Time PCR Detection System (Bio-Rad). The melting temperatures were analyzed using the derivative plots of the melting curve.

### NMR spectroscopy

NMR experiments were acquired on Bruker 800-MHz spectrometers (Bruker BioSpin, Karlsruhe, Germany). The assignments of NN206 backbone ^15^N, ^1^H^N^, ^13^Cα, ^13^Cβ, and ^13^C′ chemical shifts were obtained by ^15^N-^1^H HSQC, HNCACB, CBCA(CO)NH, HNCA, HN(CO)CA and HNCO spectra^36^. NMR samples containing 50 μM [U-15N]-labeled NN206 in the absence and presence of 100 μM Ig2 are prepared in the buffer: 50 mM sodium phosphate (pH 6.5) with 10% D2O. For temperature cycling experiments, the first 2D ^1^H–^15^N TROSY HSQC spectrum was acquired at 32 °C; the same sample was heated to 55 °C for recording the second spectra, and the temperature is decreased to 32 °C for recording the third spectra. For characterization of substrate conformation selection by NTD, 2D ^1^H-^15^N TROSY HSQC NMR spectra of 50 μM [U-^15^N]-labeled Ig2D5 titrated with 50 µM unlabeled NN206, MtaLonA or AAAP proteins were recorded during a thermal cycle starting from 32 to 60 °C, and then back to 32 °C. 72 residues of Ig2D5 were selected for calculating folded ratio. The interaction of NN206 with Sul20 peptide was recorded with 50 μM [U-15N]-labeled NN206 titrated with unlabeled 50 and 150 μM unlabeled Sul20 peptide. The interaction of NN206 with TCEP-treated denatured lysozyme was recorded with 50 μM [U-15N]-labeled NN206 titrated with 25 μM unlabeled denatured lysozyme. ^1^H-^15^N NOE data were recorded in an interleaved manner: one spectrum with 4 s recycle delay followed by 4 s saturation and another spectrum with no saturation and 8 s recycle delay. All NMR data were processed and analyzed by XWIN-NMR (Bruker BioSpin), NMRPipe^37^, and NMRView^38^.

## Data availability

The atomic coordinates and structure factors have been deposited in the Protein Data Bank, www.pdb.org, with accession codes 7CR9 and 7CRA. Other relevant data are available from the corresponding author upon reasonable requests.

## Acknowledgements

We thank Dr. Shang-Te Danny Hsu for helpful discussion. We also thank Ms S.-L. Huang of Ministry of Science and Technology (National Taiwan University) for the assistance in NMR experiments. We are grateful to the staff of the Biomedical Resource Core at the First Core Labs, National Taiwan University College of Medicine, for technical assistance. Portions of this research were carried out at beamlines 13B1 and 13C1 of the National Synchrotron Radiation Research Center (Taiwan). This work was supported by National Taiwan University Grants 108L7809 and 109L7809; and the Ministry of Science and Technology, Taiwan (grant numbers: MOST108-2113-M-002-006- and MOST109-2113-M-002-018- to S.-R.T. and MOST105-2320-B-001-015-MY3 to C.-I.C.).

## Author contributions

S.R.T., and C.I.C. conceived the study. S.R.T., and C.I.C. designed the experiments and analyzed the data. Y.C.T., C.C.L., Y.T.K., and S.J.H. performed the experiments. S.R.T., and C.I.C. wrote the manuscript with inputs from all other authors.

## References

1. Charette, M.F., Henderson, G.W. & Markovitz, A. ATP hydrolysis-dependent protease activity of the lon (capR) protein of Escherichia coli K-12. Proc Natl Acad Sci U S A 78, 4728–32 (1981).

2. Park, S.C. et al. Oligomeric structure of the ATP-dependent protease La (Lon) of Escherichia coli. Mol Cells 21, 129–34 (2006).

3. Goldberg, A.L., Moerschell, R.P., Chung, C.H. & Maurizi, M.R. ATP-dependent protease La (lon) from Escherichia coli. Methods Enzymol 244, 350–75 (1994).

4. Rotanova, T.V. et al. Classification of ATP-dependent proteases Lon and comparison of the active sites of their proteolytic domains. European Journal of Biochemistry 271, 4865–4871 (2004).

5. Botos, I. et al. The catalytic domain of Escherichia coli Lon protease has a unique fold and a Ser-Lys dyad in the active site. Journal of Biological Chemistry 279, 8140–8148 (2004).

6. Higashitani, A., Ishii, Y., Kato, Y. & Koriuchi, K. Functional dissection of a cell-division inhibitor, SulA, of Escherichia coli and its negative regulation by Lon. Mol Gen Genet 254, 351–7 (1997).

7. Wohlever, M.L., Baker, T.A. & Sauer, R.T. Roles of the N domain of the AAA+ Lon protease in substrate recognition, allosteric regulation and chaperone activity. Mol Microbiol 91, 66–78 (2014).

8. Ebel, W., Skinner, M.M., Dierksen, K.P., Scott, J.M. & Trempy, J.E. A conserved domain in Escherichia coli Lon protease is involved in substrate discriminator activity. J Bacteriol 181, 2236–43 (1999).

9. Roudiak, S.G. & Shrader, T.E. Functional role of the N-terminal region of the Lon protease from Mycobacterium smegmatis. Biochemistry 37, 11255–63 (1998).

10. Rudyak, S.G. & Shrader, T.E. Polypeptide stimulators of the Ms-Lon protease. Protein Sci 9, 1810–7 (2000).

11. Iyer, L.M., Leipe, D.D., Koonin, E.V. & Aravind, L. Evolutionary history and higher order classification of AAA+ ATPases. J Struct Biol 146, 11–31 (2004).

12. Melnikov, E.E. et al. Limited proteolysis of E. coli ATP-dependent protease Lon - a unified view of the subunit architecture and characterization of isolated enzyme fragments. Acta Biochim Pol 55, 281–96 (2008).

13. Adam, C. et al. Biological roles of the Podospora anserina mitochondrial Lon protease and the importance of its N-domain. PLoS One 7, e38138 (2012).

14. Cheng, I. et al. Identification of a Region in the N-Terminus of Escherichia coli Lon That Affects ATPase, Substrate Translocation and Proteolytic Activity. Journal of Molecular Biology 418, 208–225 (2012).

15. Gottesman, S. Proteolysis in bacterial regulatory circuits. Annu Rev Cell Dev Biol 19, 565–87 (2003).

16. Baker, T.A. & Sauer, R.T. ATP-dependent proteases of bacteria: recognition logic and operating principles. Trends Biochem Sci 31, 647–53 (2006).

17. Lin, C.C. et al. Structural Insights into the Allosteric Operation of the Lon AAA+ Protease. Structure 24, 667–675 (2016).

18. Su, S.C. et al. Structural Basis for the Magnesium-Dependent Activation and Hexamerization of the Lon AAA+ Protease. Structure 24, 676–686 (2016).

19. Li, M. et al. Crystal structure of the N-terminal domain of E. coli Lon protease. Protein Sci 14, 2895–900 (2005).

20. Duman, R.E. & Lowe, J. Crystal structures of Bacillus subtilis Lon protease. J Mol Biol 401, 653–70 (2010).

21. Li, M. et al. Structure of the N-terminal fragment of Escherichia coli Lon protease. Acta Crystallogr D Biol Crystallogr 66, 865–73 (2010).

22. Chen, X. et al. Crystal structure of the N domain of Lon protease from Mycobacterium avium complex. Protein Sci 28, 1720–1726 (2019).

23. Hsu, S.T., Cabrita, L.D., Fucini, P., Dobson, C.M. & Christodoulou, J. Structure, dynamics and folding of an immunoglobulin domain of the gelation factor (ABP-120) from Dictyostelium discoideum. J Mol Biol 388, 865–79 (2009).

24. Van Melderen, L. et al. ATP-dependent degradation of CcdA by Lon protease. Effects of secondary structure and heterologous subunit interactions. J Biol Chem 271, 27730–8 (1996).

25. Yang, M., Dutta, C. & Tiwari, A. Disulfide-bond scrambling promotes amorphous aggregates in lysozyme and bovine serum albumin. J Phys Chem B 119, 3969–81 (2015).

26. Gur, E. & Sauer, R.T. Recognition of misfolded proteins by Lon, a AAA(+) protease. Genes Dev 22, 2267–77 (2008).

27. Patterson, J. et al. Correlation of an adenine-specific conformational change with the ATP-dependent peptidase activity of Escherichia coli Lon. Biochemistry 43, 7432–42 (2004).

28. Vasilyeva, O.V., Kolygo, K.B., Leonova, Y.F., Potapenko, N.A. & Ovchinnikova, T.V. Domain structure and ATP-induced conformational changes in Escherichia coli protease Lon revealed by limited proteolysis and autolysis. FEBS Lett 526, 66–70 (2002).

29. McCoy, A.J., Fucini, P., Noegel, A.A. & Stewart, M. Structural basis for dimerization of the Dictyostelium gelation factor (ABP120) rod. Nat Struct Biol 6, 836–41 (1999).

30. Wernimont, A. & Edwards, A. In situ proteolysis to generate crystals for structure determination: an update. PLoS One 4, e5094 (2009).

31. Otwinowski, Z. & Minor, W. [20] Processing of X-ray diffraction data collected in oscillation mode. Methods Enzymol 276, 307–326 (1997).

32. McCoy, A.J. Solving structures of protein complexes by molecular replacement with Phaser. Acta Crystallogr D Biol Crystallogr 63, 32–41 (2007).

33. Terwilliger, T.C. et al. Iterative model building, structure refinement and density modification with the PHENIX AutoBuild wizard. Acta Crystallogr D Biol Crystallogr 64, 61–9 (2008).

34. Emsley, P. & Cowtan, K. Coot: model-building tools for molecular graphics. Acta Crystallogr D Biol Crystallogr 60, 2126–32 (2004).

35. Murshudov, G.N. et al. REFMAC5 for the refinement of macromolecular crystal structures. Acta Crystallogr D Biol Crystallogr 67, 355–67 (2011).

36. Sattler M, S.J., Griesinger C. Heteronuclear multidimensional NMR experiments for the structure determination of proteins in solution employing pulsed field gradients. Progress in Nuclear Magnetic Resonance Spectroscopy. 34, 93–158 (1999).

37. Delaglio, F. et al. NMRPipe: a multidimensional spectral processing system based on UNIX pipes. J Biomol NMR 6, 277–93 (1995).

38. Downing, A.K. Protein NMR techniques, xi, 487 p. (Humana Press, Totowa, N.J., 2004).

