## Supplementary Figs for "Molecular insights into substrate recognition and discrimination by the N-terminal domain of Lon AAA+ protease"

**This PDF file includes:**

Supplementary Figs. S1 to S8

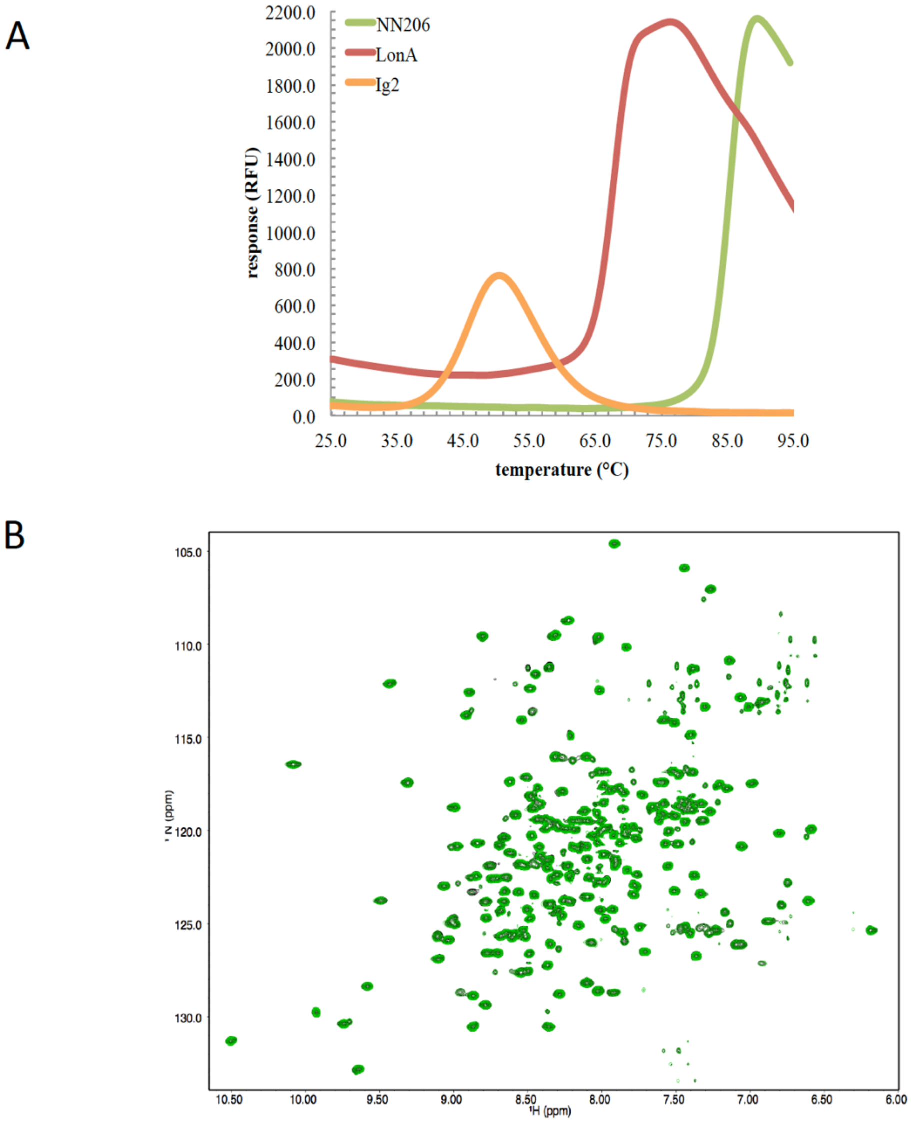


Fig. S1. ***The stability of MtaLonA, NTD and substrate Ig2.*** (A) SYPRO Orange is used to monitor the availability of hydrophobic areas of thermally unfolded protein in the thermal shift assay. The melting temperatures of substrate Ig2 is 46 °C, suggesting that the structure of Ig2 is thermal denatured as a substrate of MtaLonA at 55 °C. The $T_m$ of MtaLonA NTD and full-length MtaLonA are 85.5 and 68.0 °C, respectively. (B) MtaLonA NTD remains stable over the entire thermal cycling. 2D $^{15}N$-$^{1}H$ TROSY HSQC NMR spectra of apo NN206 are overlaid and compared before (black) and after (green) one thermal cycle.

A

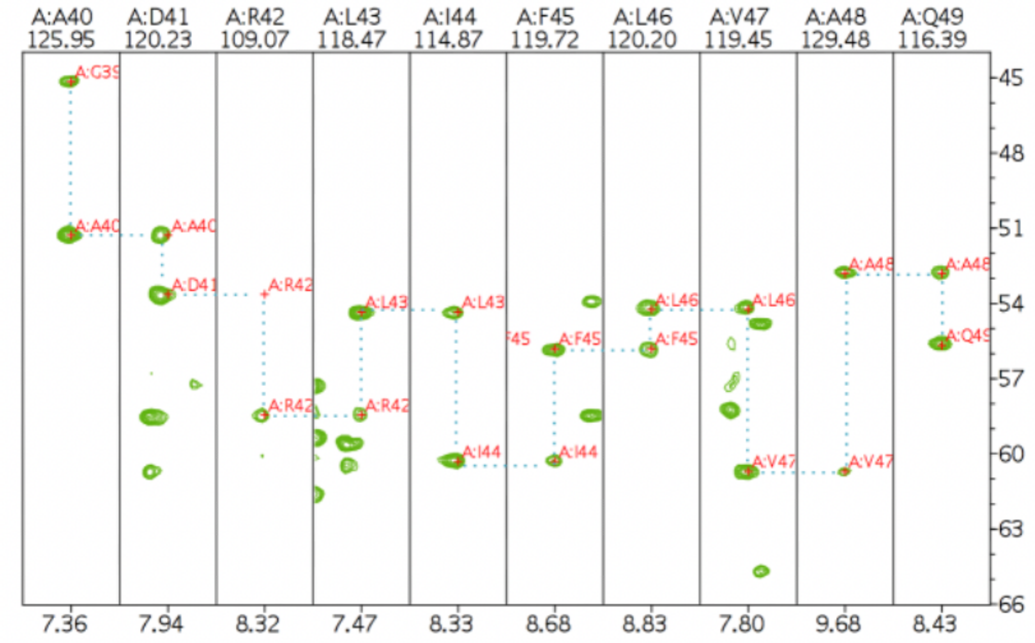


B

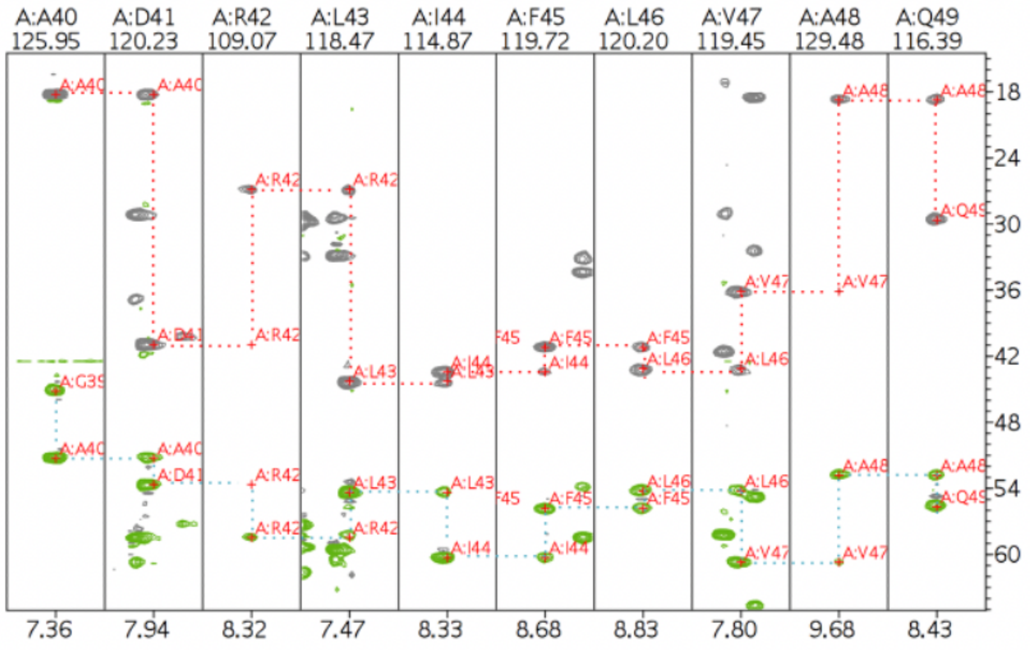


Fig. S2***. Backbone amide resonances of NN206.*** Strip plots of HNCACB (A) and HNCACB (B) spectra and the sequential connectivity is shown from A40 to Q49.

*Figure S2 continued on next page*

C

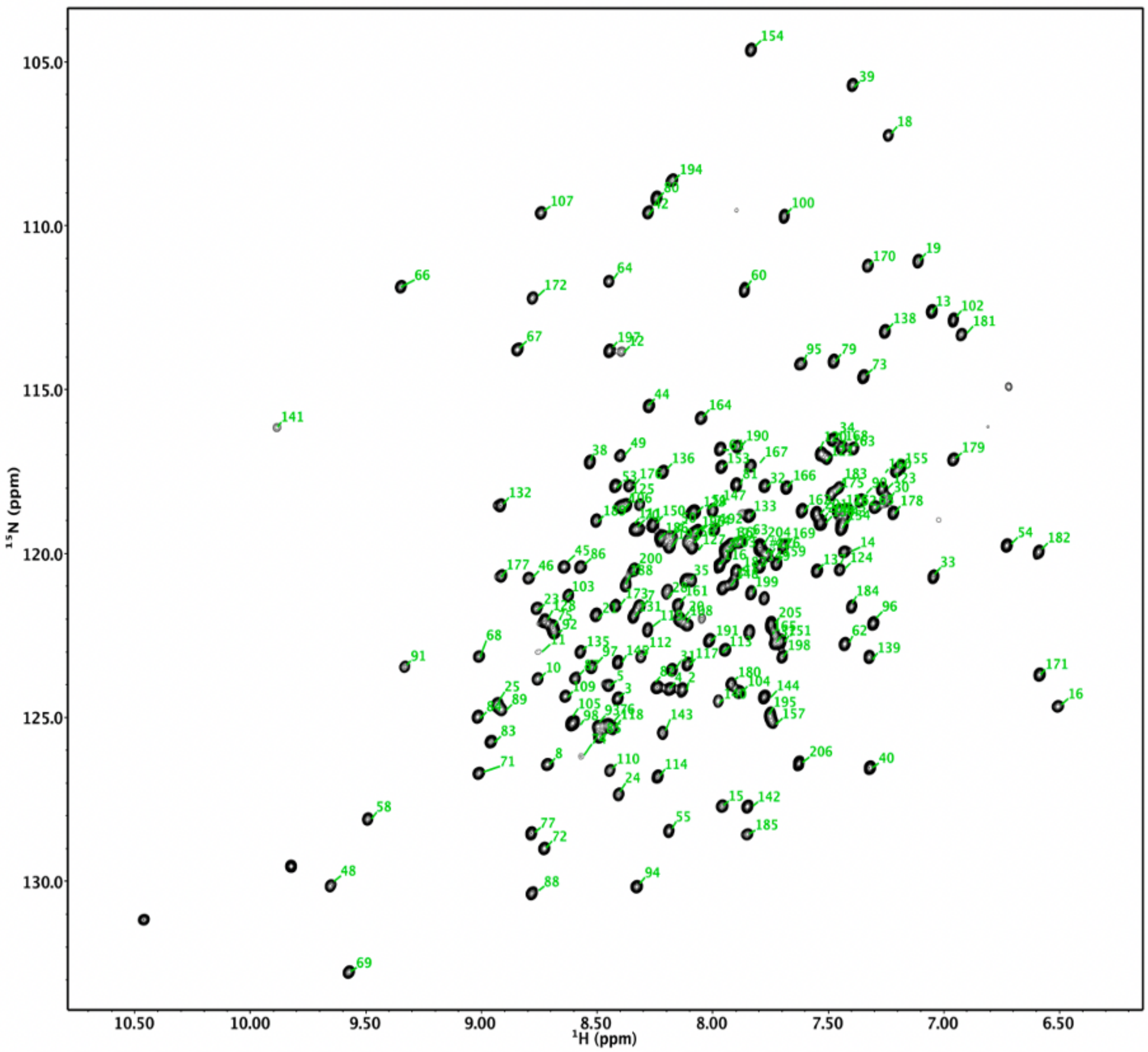


*Figure S2 continued*

(C) $^{15}N$-$^{1}H$ TROSY HSQC spectrum of NN206: backbone amide resonances are labeled according to residue number.

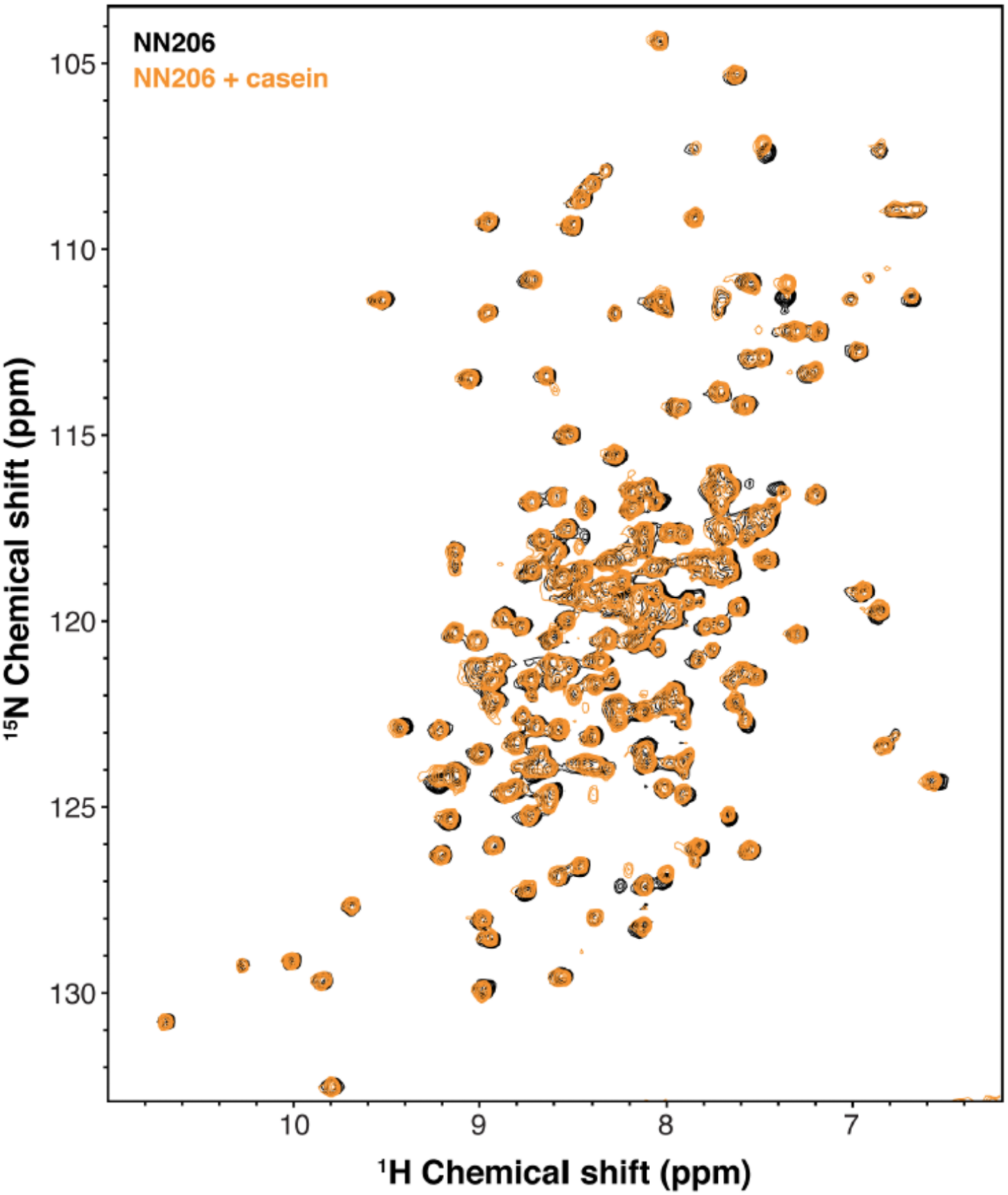


Fig. S3. Overlay of the $^{15}$N-$^{1}$H HSQC spectra of NN206 in apo (black) and casein-bound (orange) states recorded at 55 °C.

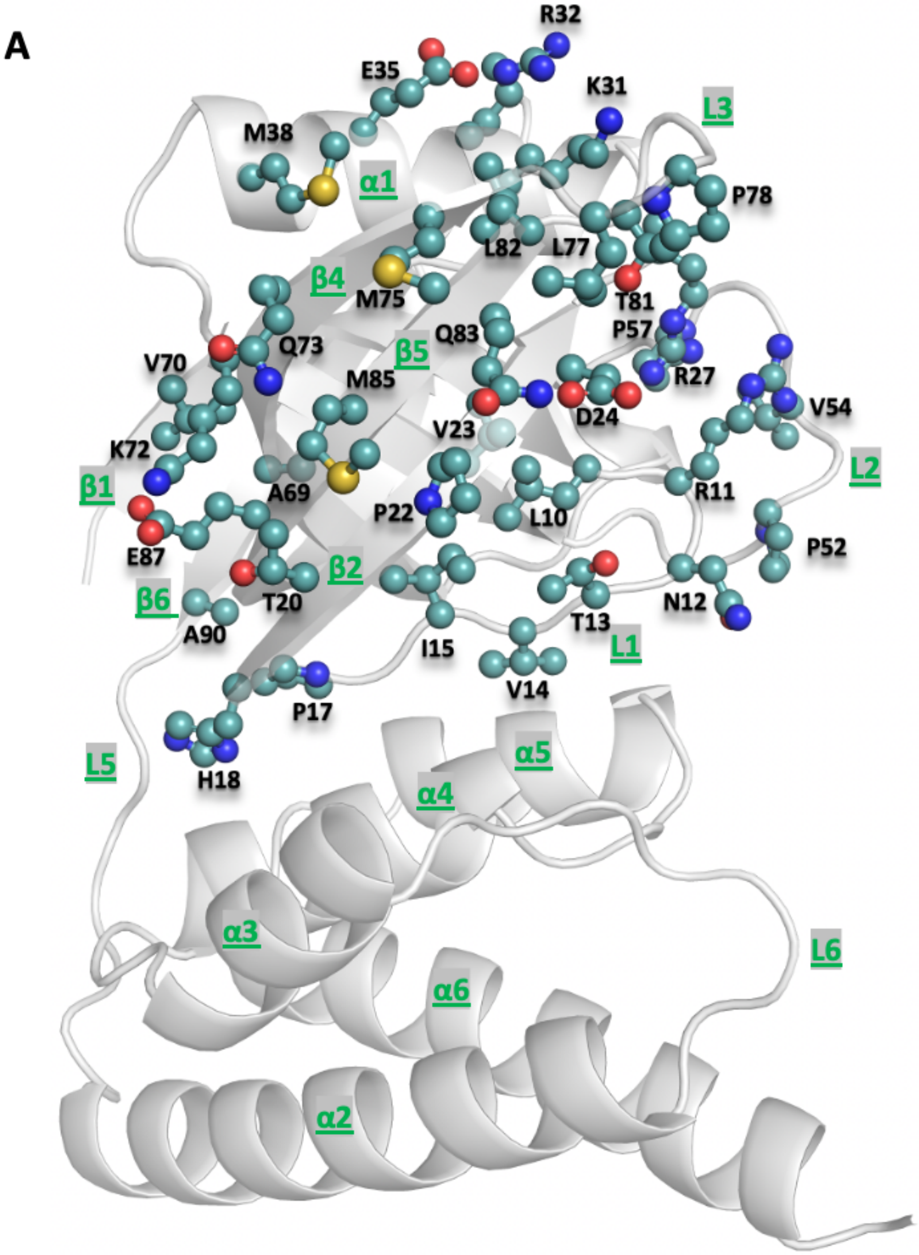


Fig. S4. ***NMR analysis indicated that thermally damaged Ig2 induced significant CSPs and broadened resonances in the NTD N-lobe.*** (A) Thermally damaged Ig2 induced significant CSPs and broadened resonance mainly at helix α1, loop L1, L2, L3 and β-sheet β2/β5/β4, which consist primarily of hydrophobic residues and are decorated by a number of polar residues.

*Figure S4 continued on next page*

B

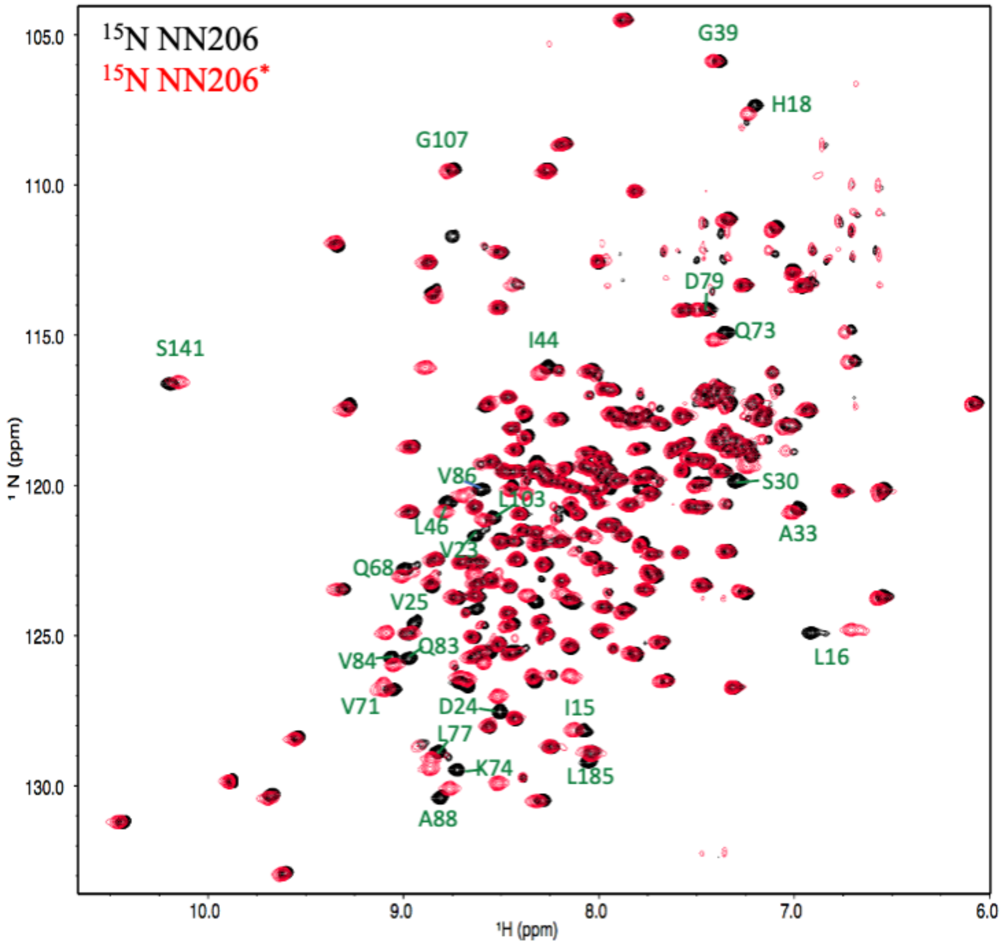


*Figure S4 continued*

(B) Overlay of the $^{15}$N-$^{1}$H HSQC spectra of apo NN206 (black) and NN206* (red) states recorded at 55 °C. Spectral analysis shows chemical shift differences between NN206 and NN206* are negligible, suggesting that they have very similar structures, with the exception of the region surrounding the point substitutions.

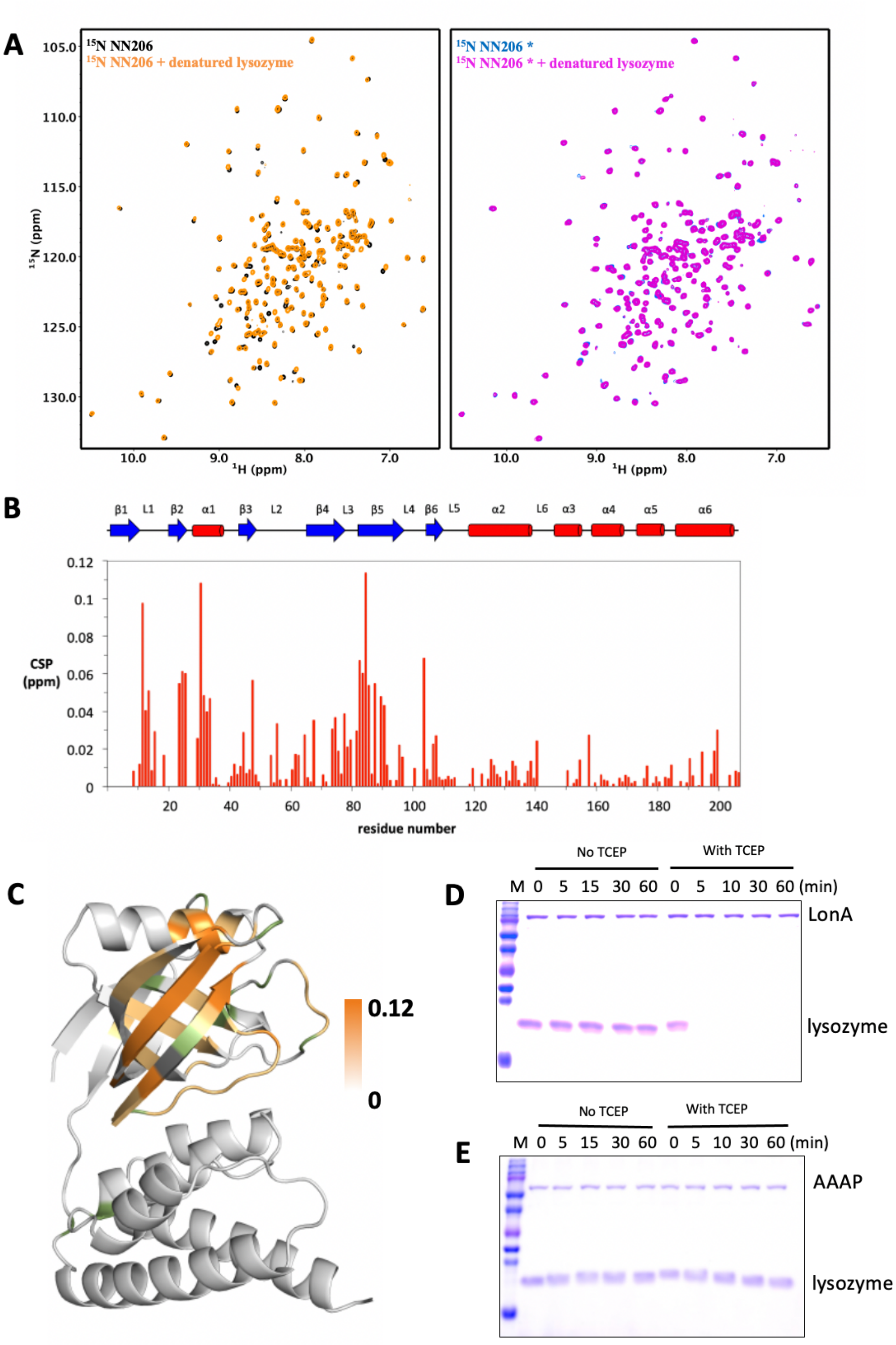


Fig. S5 ***Interaction of NTDs with the scrambled lysozyme.***

(A) Overlay of $^{1}$H–$^{15}$N HSQC spectra of 50 μM NN206 in the absence (black) and presence (orange) of 25 μM TCEP-treated denatured lysozyme. Overlay of $^{1}$H–$^{15}$N

HSQC spectra of 50 μM NN206* in the absence (blue) and presence (magenta) of TCEP-treated denatured 25 μM lysozyme. (B) CSPs of amide moieties of 50 μM NN206 after binding 25 μM, plotted against the NN206 amino acid residue number. The secondary structure of the native-state NN206 is indicated by the blue bar on the top of the chart. (C) Structural mapping of the chemical shift perturbations (orange) of NN206 caused by TCEP-treated denatured lysozyme. 12 proline residues are shown in green. (D) Degradation of native and denatured lysozyme by full-length MtaLonA. (E) Degradation of native and denatured lysozyme by AAAP.

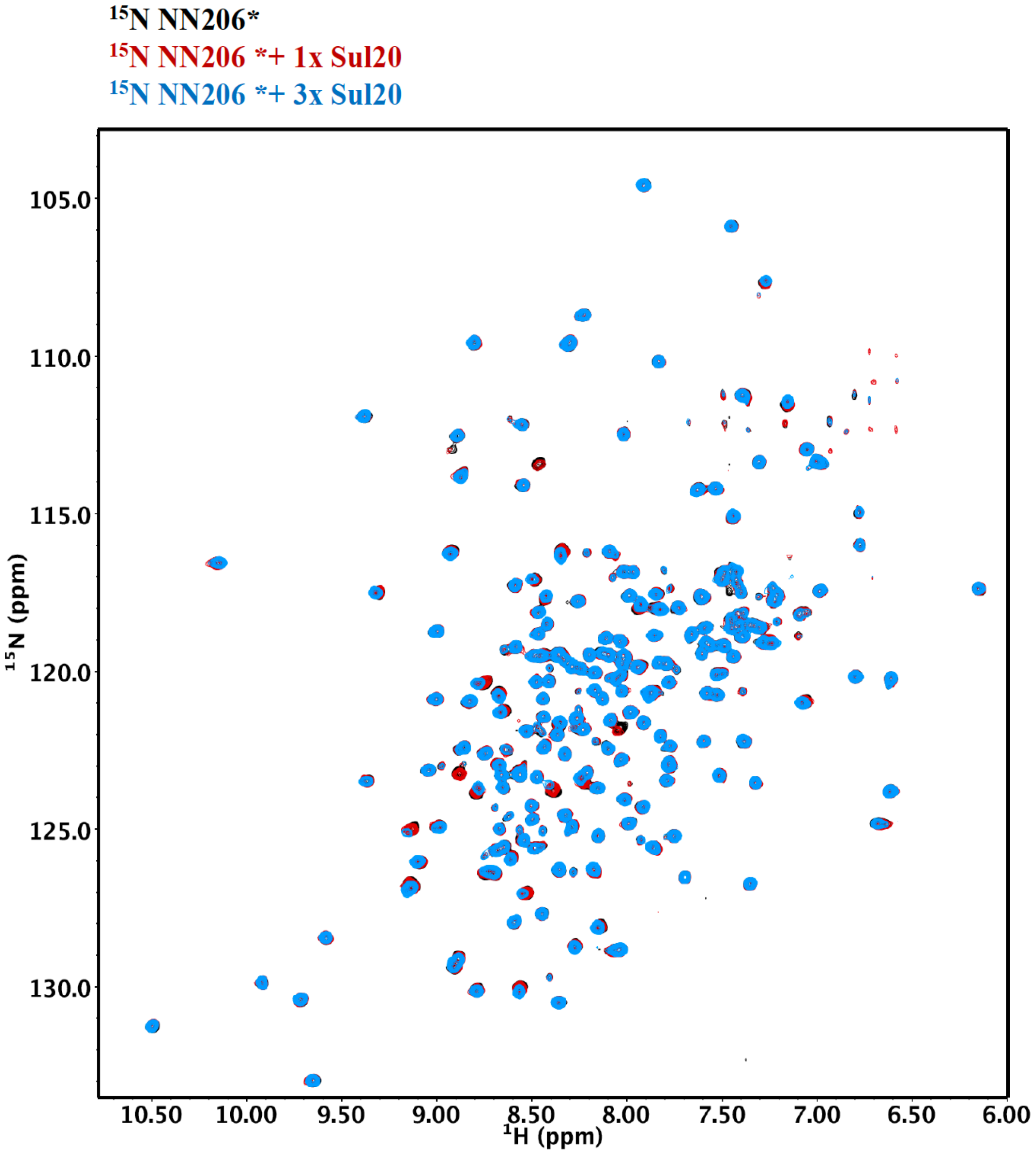


Fig. S6 ***Interaction of MtaLonA NTD* with degron tag Sul20.***

(A) Two-dimensional $^{15}$N-$^{1}$H HSQC spectra of 50 μM NN206* titrated with increasing concentration of Sul20 peptide: 0 (black), 50 (red), and 150 μM (blue).

*Figure S6 continued on next page*

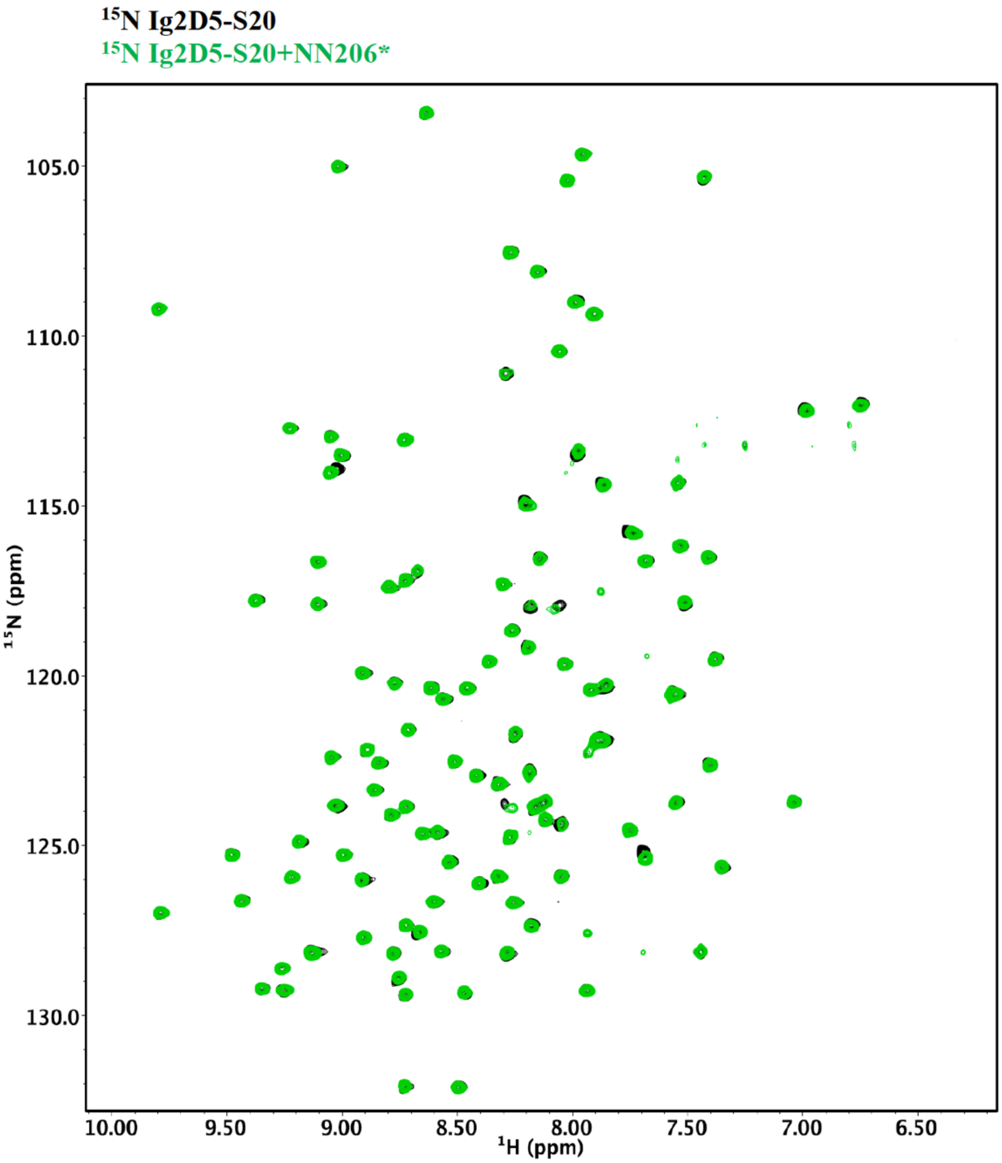


*Figure S6 continued*

(B) $^{1}$H-$^{15}$N NMR spectra of Ig2D5-S20 in the absence (black) and in the presence (green) of an equimolar amount of NN206*.

A

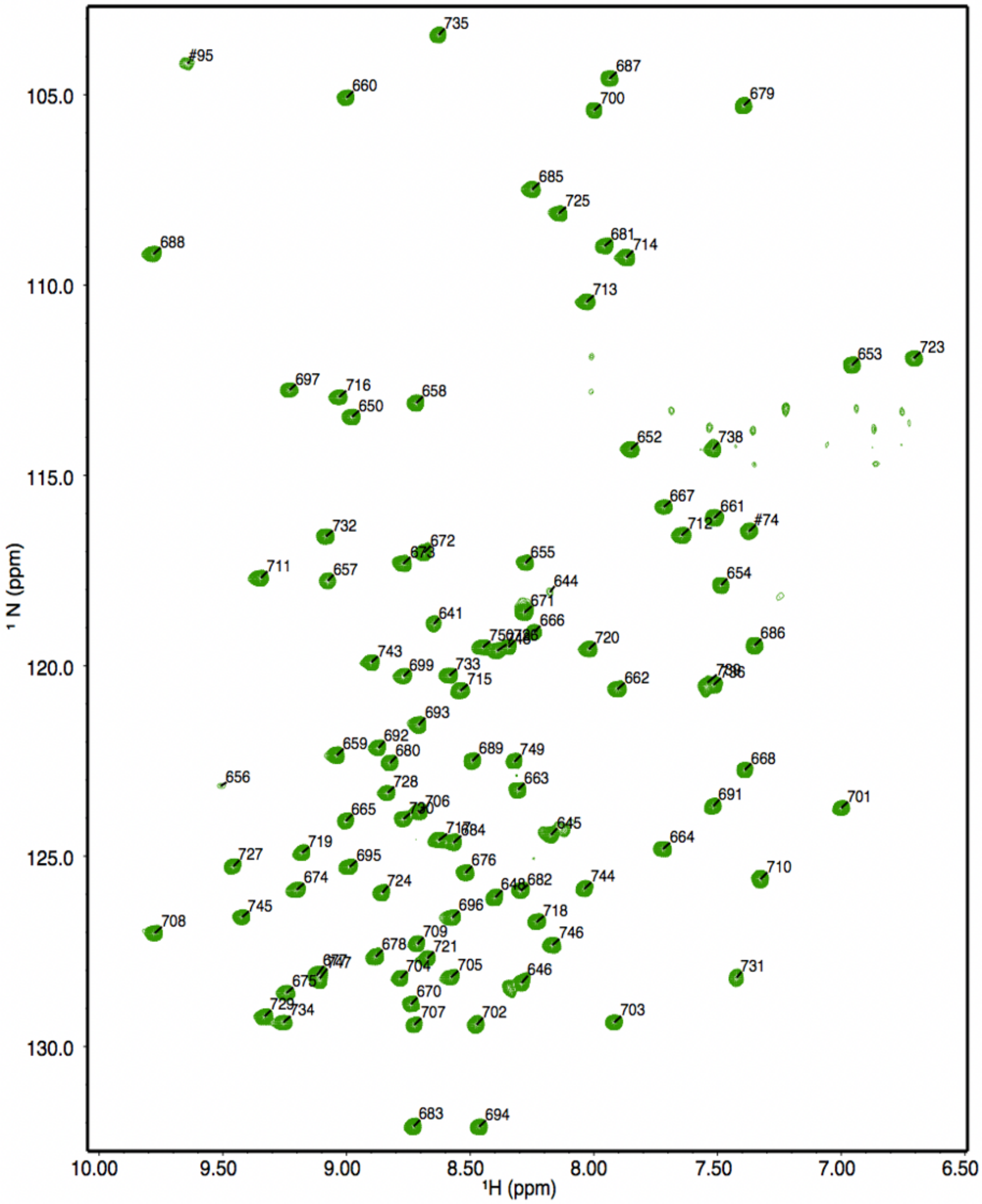


Fig. S7. ***The thermodynamics of Ig2D5 unfolding-refolding is in the presence of NN206*.*** (A) 2D $^{1}$H-$^{15}$N TROSY HSQC NMR spectrum of Ig2D5 (green): backbone amide resonances are labeled according to residue number and 72 residues are selected for calculating folded ratio.

*Figure S7 continued on next page*

B

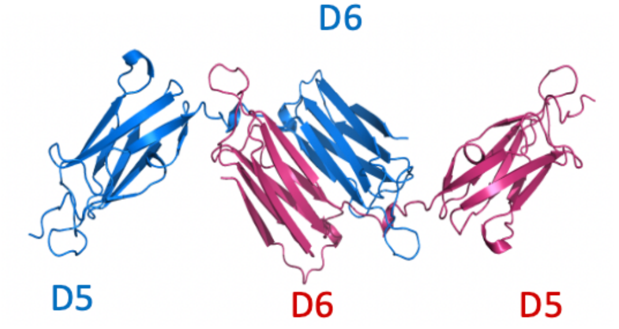


C

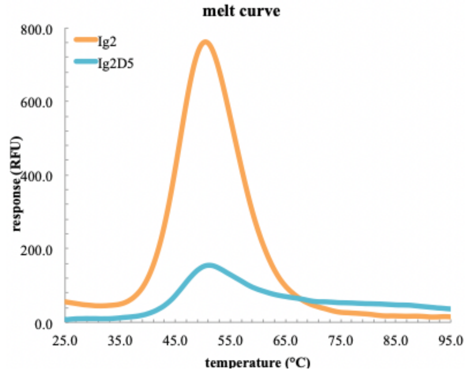


D

E

F

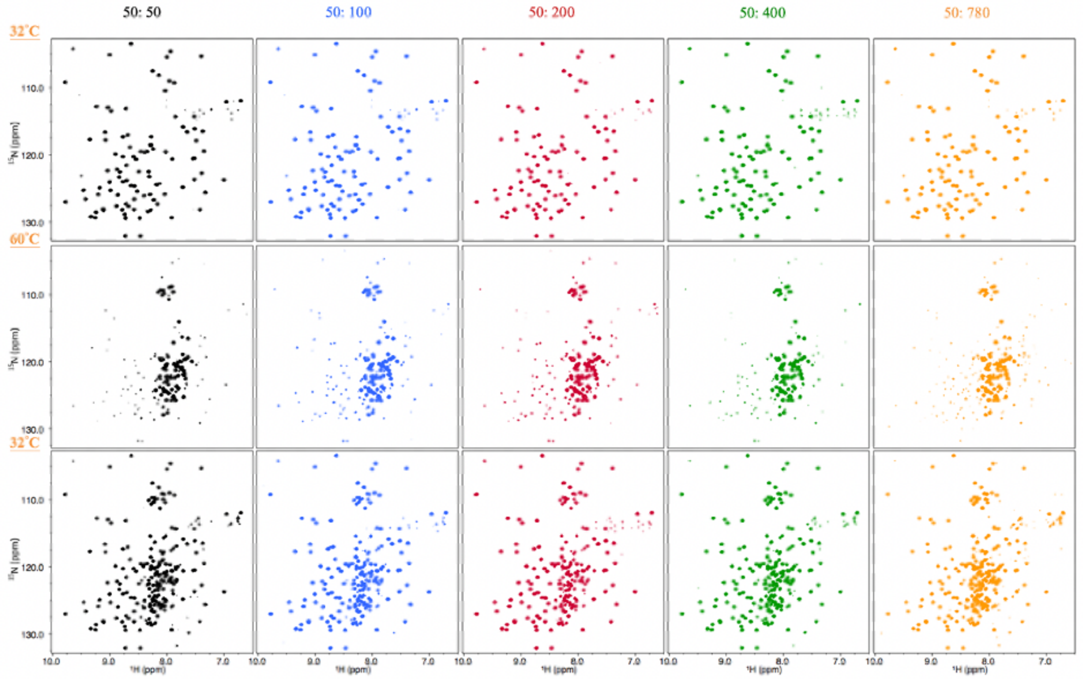


(B) domains 5 and 6 of the gelation factor ABP-120 from *Dictyostelium discoideum* (PDB code:1QFH). (C) The $T_m$ of substrate Ig2D5 is similar to Ig2. (D) 2D $^{1}H$-$^{15}N$ TROSY HSQC NMR spectra of Ig2D5 titrated with 50 (black), 100 (blue), 200 (red), 400

(green) or 780 (orange) μM NN206* are recorded during a thermal cycle starting from 32 (D) to 60 °C (E), and then returning to 32 °C (F).

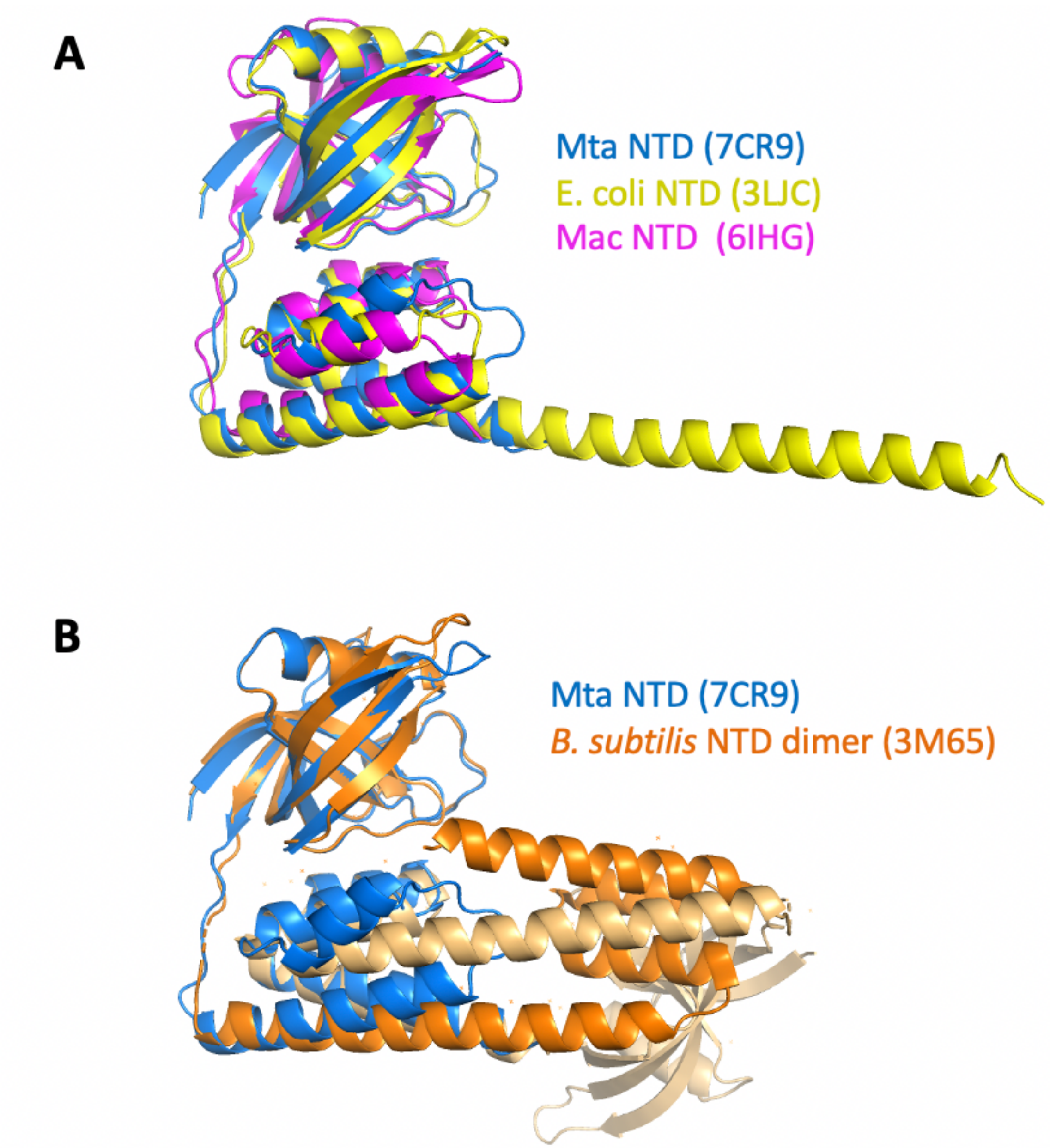


Fig. S8. ***Structural superposition of the N-terminal fragments from MtaLon, EcLon, MacLon and BsLon.*** (A) The structures of the N-terminal fragments from MtaLon, EcLon and MacLon showed that the N- and the C-lobes are joined together via a short linker. The PDB codes are indicated in parentheses. (B) The N-terminal fragment from *B. subtilis* LonA was previously reported to adopt a domain-swapped dimer where the N-lobe of one monomer is positioned next to the C-lobe of the other monomer.

*Figure S8 continued on next page*

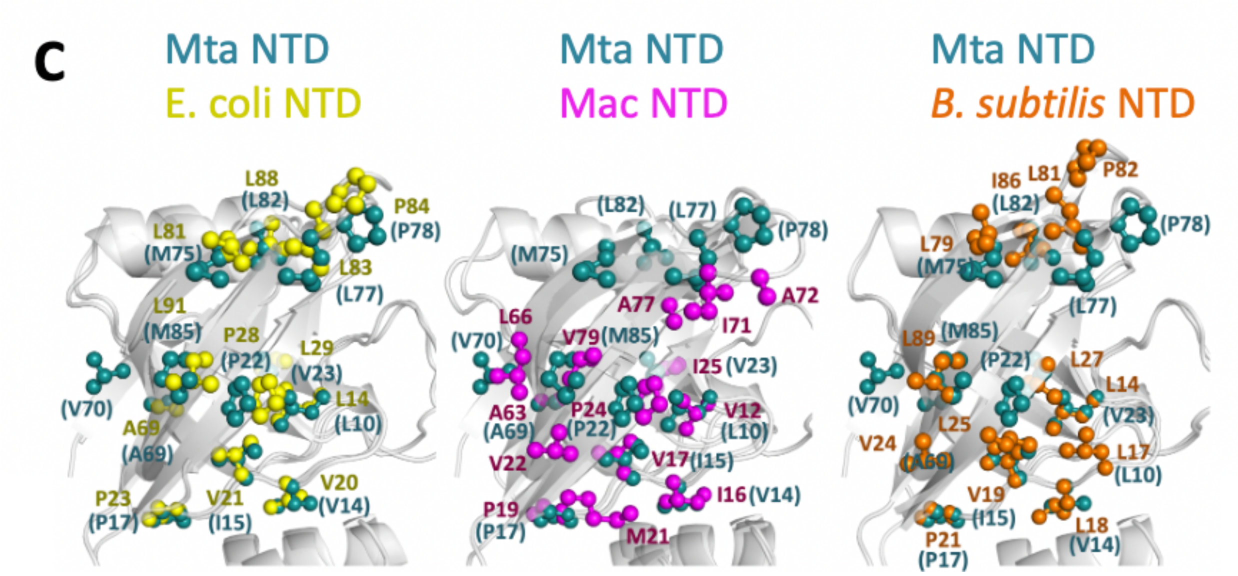


*Figure S8 continued*

(C) All four structures exhibit highly similar exposed hydrophobic residues at the N-lobes of their NTD.
